# Extrachromosomal circular DNA promotes inflammation and hepatocellular carcinoma development

**DOI:** 10.1101/2025.01.22.634016

**Authors:** Lap Kwan Chan, Juanjuan Shan, Elias Rodriguez-Fos, Marc Eamonn Healy, Peter Leary, Rossella Parrotta, Nina Desboeufs, Gabriel Semere, Nadine Wittstruck, Anton G. Henssen, Achim Weber

## Abstract

Two decades after the initial report on increased micronuclei in human chronic liver disease (CLD) and hepatocellular carcinoma (HCC), their role in HCC development is still poorly understood. Here we show that micronuclei in hepatocytes trigger hepatic immune response and promote HCC development via an increased level of extrachromosomal circular DNA (eccDNA). Livers from a CLD model (*Mcl1*^Δhep^ mice) show increased micronuclei in parallel to eccDNA. Circular sequencing confirms higher eccDNA levels in micronuclei compared to primary nuclei. We developed nuclei-segregated DNA fiber (NuSeF) assay to show that micronuclei are more susceptible to replication stress, showing increased replication fork slowing. By comparing different murine liver disease models, high eccDNA is correlated with increased tumor incidence. EccDNA is a strong immunostimulant and promotes a crosstalk between hepatocytes and immune cells through the cGAS-STING pathway. Deletion of *Sting1* in *Mcl1*^Δhep^ mice reduces immune cell chemotaxis, as well as tumor incidence. Our findings suggest that eccDNA from micronuclei mediates inflammation-driven liver carcinogenesis in CLD.

## Introduction

In 2020, primary liver cancer ranked as the sixth most common type of cancer and the third leading cause of cancer-related deaths (1). The number of new diagnoses is expected to increase over the next two decades, reaching 1.4 million in 2040. Hepatocellular carcinoma (HCC) is the most prevalent type of liver cancer and accounts for over 80% of all primary liver cancers (2). Most HCCs develop following a long-term chronic liver disease (CLD) and, in recent years, increased incidence caused by metabolic dysfunction-associated steatotic liver diseases (MASLD). Chronic liver inflammation and fibrosis, which are associated with constant liver cell death, can subsequently progress to cirrhosis and significantly increase the risk of HCC development. Hepatocyte hyperproliferation is a phenomenon frequently observed in CLD as a compensatory regenerative mechanism triggered by increased cell death and supported by a proinflammatory microenvironment (3, 4). The production of immunogenic molecules, known as damage-associated molecular patterns (DAMPs), from damaged cells is responsible for triggering immune responses and sustains the status of ‘sterile inflammation’ (5). Under such proinflammatory microenvironments, hepatocytes can experience increased DNA damage and genetic instability, which can eventually promote liver carcinogenesis (6).

Myeloid cell leukemia 1 (MCL1) is a pro-survival member of the B-cell lymphoma 2 (Bcl-2) family which shares BH domains with Bcl-2 and Bcl-X_L_ (7). MCL1 inhibits apoptosis by sequestering the pro-apoptotic proteins Bcl-2 homologous antagonist killer (Bak) and Bcl-2-associated protein X (Bax), which prevents mitochondrial outer membrane permeabilization (MOMP) during the initiation of apoptosis. Our group previously reported that liver-specific knockout of *Mcl1* in mice (*Mcl1*^Δhep^) resulted in a phenotype resembling CLD in patients characterized by elevated apoptosis, compensatory proliferation, and DNA damage, with increased tumor incidence at 12 months (4, 6). Particularly, the increase in DNA damage and genomic instability in hepatocytes may have a direct link to carcinogenesis. However, it is still unclear if the carcinogenesis from CLD to HCC is mediated by a cell-autonomous or non-cell-autonomous mechanism. In the current study, we used *Mcl1*^Δhep^ mice as a CLD model and discovered a novel mechanism involving a crosstalk between hepatocytes and immune cells in the liver, mediated by extrachromosomal circular DNA (eccDNA), driving a proinflammatory microenvironment in a manner that was dependent on the cGAS-STING pathway. Our findings suggested that a non-cell-autonomous mechanism was involved in liver tumorigenesis, with micronuclei and eccDNA playing a pivotal role. Moreover, we developed a new method that enabled us to study replication stress within primary nuclei and micronuclei, and therefore provided mechanistic insights into eccDNA generation.

## Results

### Livers of *Mcl1*^Δhep^ mice show increased micronuclei and activated cGAS-STING pathway

Previously, we reported that *Mcl1*^Δhep^ mice already exhibited liver hyperproliferation at 2 months of age (4, 6). To determine if the observed increase in DNA damage in *Mcl1*^Δhep^ livers was a consequence of increased replication stress, we performed DNA fiber assay on primary hepatocytes from wild-type (WT) and *Mcl1*^Δhep^ mice. We established MCL1 immunohistochemistry to confirm a high knockout efficiency and observed that MCL-1 expression was restricted to non-parenchymal cells (NPC) in *Mcl1*^Δhep^ livers (Fig. 1A). Pulse labeling of two thymidine analogs, chlorodeoxyuridine (CldU) and iododeoxyuridine (IdU), allowed monitoring of the replication fork dynamics and detection of replication stress in cells (8, 9). We observed that MCL1-deficient hepatocytes displayed a significantly longer CldU tract length with no difference in the total tract length (CldU+IdU) (Fig. 1B & Fig. S1A). The ratio of IdU to CldU, however, was significantly reduced in *Mcl1*^Δhep^ hepatocytes compared to WT (Fig. 1C). This suggested that *Mcl1*^Δhep^ hepatocytes had a higher rate of replication fork stalling, a sign of increased replication stress that was detected during IdU labeling. Since the formation of micronuclei had been associated with increased DNA damage and replication stress (10), we examined the level of micronucleated hepatocytes in *Mcl1*^Δhep^ livers. Using Δ-catenin as a membrane marker, we visualized and founda significantly higher level of micronucleated hepatocytes in the *Mcl1*^Δhep^ compared to WT livers (Fig. 1D&E, Fig. S1B). These results suggested that MCL1 deficiency in hepatocytes caused a higher replication stress and led to increased micronuclei formation.

**Fig. 1.**
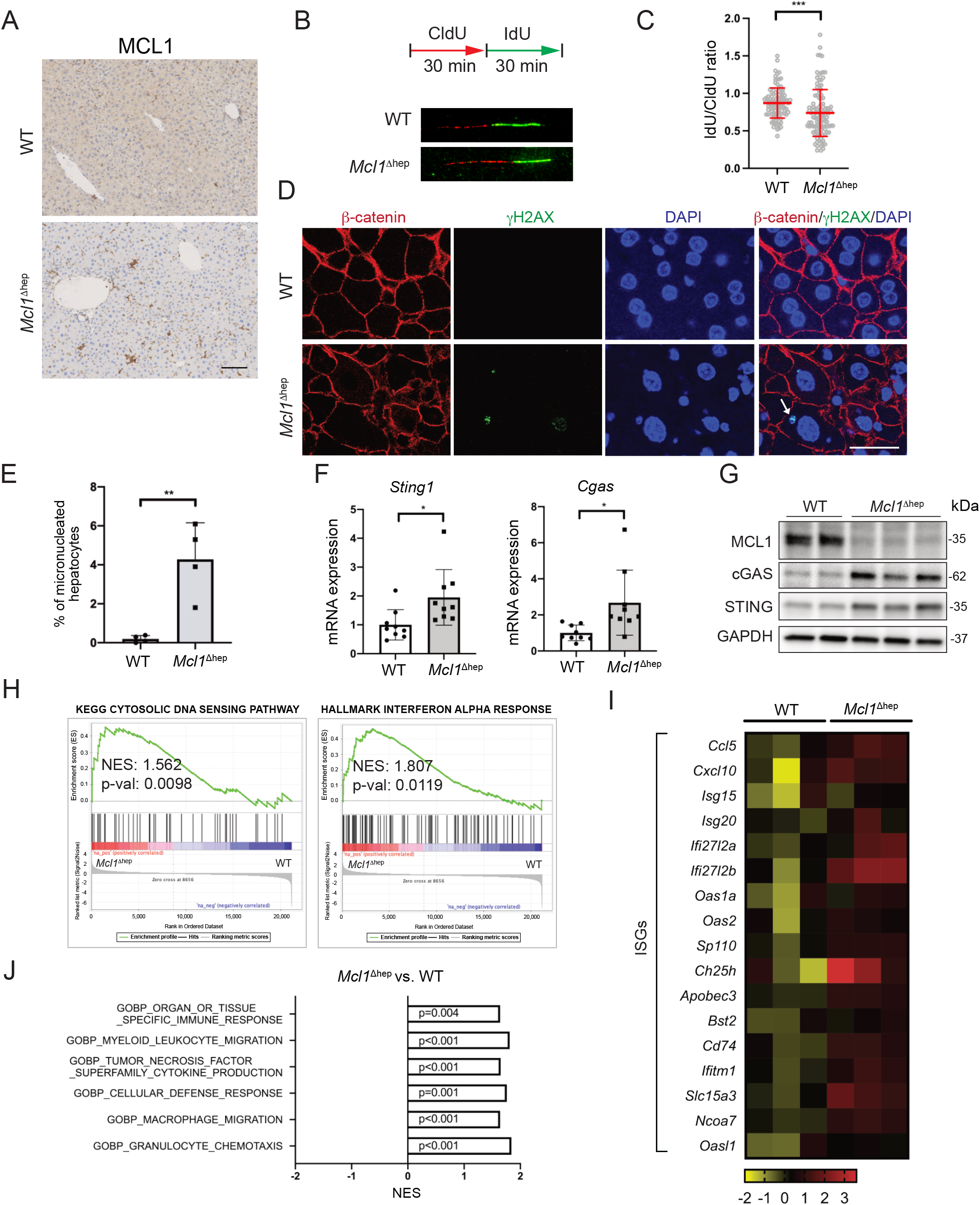
Deletion of MCL1 in hepatocytes leads to increased micronucleated hepatocytes and an activation of the cGAS-STING pathway. A) MCL1 IHC staining on WT and *Mcl1*^Δhep^ livers. Scale bar: 100μm. B) DNA fiber assay on primary hepatocytes from WT and *Mcl1*^Δhep^ mice after 24 hours in culture. CldU: 5-chloro-2’deoxyuridine; IdU: 5-Iodo-2’-Deoxyuridine. C) Quantification of IdU to CldU ratio. D) Immunofluorescence staining of 2-month-old liver tissue showing the presence of micronuclei (arrow) in hepatocytes. Scale bar: 25μm. E) Quantification of micronucleated hepatocytes as a percentage of the total hepatocytes. n=4. F) Quantitative PCR showing upregulation of the cGAS and STING expression in the *Mcl1*^Δhep^ liver. n=9. G) Immunoblot of protein lysates from 2-month-old WT and *Mcl1*^Δhep^ livers showing upregulation of the cGAS and STING protein levels. H) GSEA showing an enrichment of the cytosolic DNA sensing pathway and the interferon alpha response in *Mcl1*^Δhep^ livers. NES: net enrichment score. I) Heat-map showing upregulation of ISGs in *Mcl1*^Δhep^ livers. J) *Mcl1*^Δhep^ mice showed significant enrichment of various gene signatures involved in immune responses and immune cell recruitment. Student’s t-test ^*^ p<0.05; ^**^ p<0.01; ^***^ p<0.001.

The cytosolic DNA sensing pathway was recently reported to connect DNA damage to the activation of IRF3 and NF-κB, both known as important transcription factors for immune response (11). We, therefore, analyzed if this pathway was activated in the *Mcl1*^Δhep^ liver. The expression of the key components of this pathway, cGAS, and STING, were upregulated in 2- and 12-month-old *Mcl1*^Δhep^ livers both at mRNA and protein levels (Fig. 1F&G & Fig S1C). Transcriptomic analysis of 2-month-old liver samples showed an enrichment of the cytosolic DNA sensing pathway and the interferon alpha response signatures in the *Mcl1*^Δhep^ liver (Fig. 1H). Activation of the cGAS-STING pathway is known to facilitate the production of various interferon-stimulated genes (ISGs) (10). Analysis of an ISG gene list showed that these genes were upregulated in the 2-month-old *Mcl1*^Δhep^ liver, including chemokines (*Ccl5* and *Cxcl10*) and interferon alpha inducible proteins (*Ifi27l2a* and *Ifi27l2b*) (Fig. 1I). Several immune response-related signatures, such as pathways promoting immune cell chemotaxis, were also enriched in *Mcl1*^Δhep^ livers (Fig. 1J). To corroborate these results, we performed immunohistochemistry and found that the infiltration of B cells (B220^+^), neutrophils (Ly6G^+^), and macrophages (F4/80^+^) was significantly higher in *Mcl1*^Δhep^ livers (Fig. S2A&B).

Thus an activation of the cGAS-STING pathway was observed in *Mcl1*^Δhep^ livers, even though hepatocytes had been shown to have an incomplete cGAS-STING pathway (12-14). Via immunohistochemical staining, we observed that both hepatocytes and NPC expressed cGAS (Fig. S2C). Expression of STING, however, was restricted to NPC (Fig. S2C&D). Using serial sections, we visualized a co-localization of F4/80^+^ and STING (Fig. S2E). This pattern was further confirmed from isolated immune cells and hepatocytes using Western blot (Fig. S2F). Due to the observed differences in the spatial expression pattern of cGAS and STING, we sought to determine whether there is a mechanism mediating a crosstalk between DNA damage in hepatocytes and the activation of the cGAS-STING pathway in immune cells. One of the reported mechanisms to promote a crosstalk between cGAS-activated cells and the adjacent bystander cells is via the production and transfer of the second messenger molecule cGAMP (15, 16). Measurement of cGAMP using ELISA showed that there was no significant change in the cGAMP level between WT and *Mcl1*^Δhep^ liver lysates (Fig. S2G). Therefore, presumably, other mechanisms are involved in facilitating the crosstalk between hepatocytes and immune cells in the *Mcl1*^Δhep^ liver.

### EccDNA is increased in *Mcl1*^Δhep^ liver and have a strong immunostimulatory activity

EccDNA is a class of circular DNA elements that can be found in different tissues (17). The biogenesis of eccDNA is still not completely understood, though recent evidence showed that their generation might be related to DNA damage, increased cellular stress, and apoptosis (18, 19). We performed eccDNA isolation using a protocol established for cultured cells and adapted it for liver tissues (Fig. 2A) (20). We observed that 2-month-old *Mcl1*^Δhep^ livers contained more eccDNA compared to age-matched WT controls (Fig. 2B). Using electron microscopy, we were able to visualize and confirm the presence of eccDNA, using a plasmid (pRSV) as a control (Fig. 2C & S3A). To verify our hypothesis that the observed activation of the cGAS-STING pathway in *Mcl1*^Δhep^ livers was due to the activities in immune cells, we isolated immune cells and examined their STING and p-p65 levels. Immune cells from *Mcl1*^Δhep^ livers showed a stronger STING expression and higher NF-κB activities, as indicated by increased phosphorylation of p65 at S536 (21), compared to immune cells from WT livers (Fig. 2D&E).

**Fig. 2.**
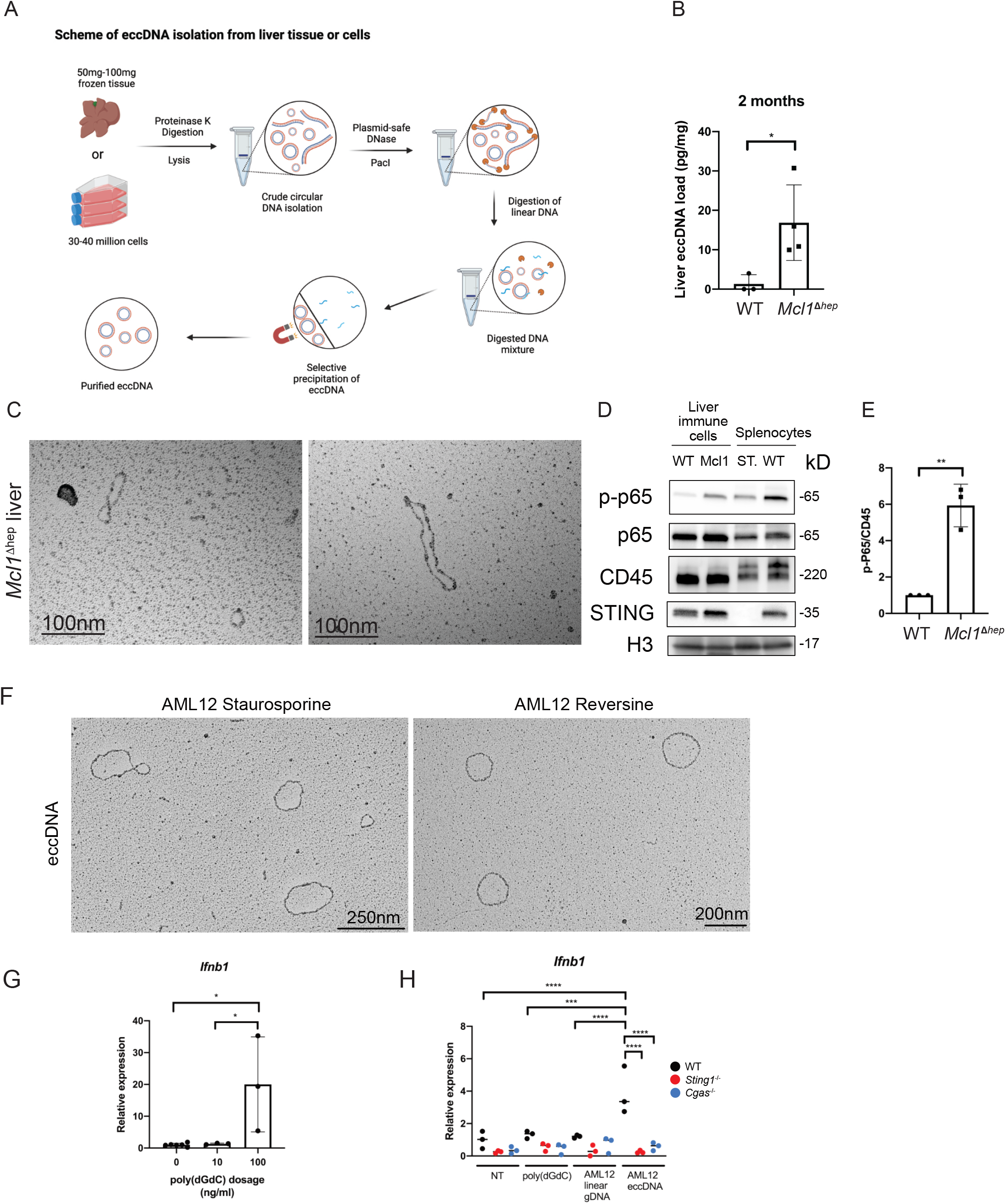
The crosstalk between MCL1-deficient hepatocytes and immune cells is through an increase in eccDNA. A) Schematic representation showing the steps of eccDNA isolation from liver tissues and cultured cells. B) Quantification of eccDNA isolated from WT and *Mcl1*^Δhep^ livers by Qubit HS dsDNA kit (pg/mg of tissues). Student’s t-test: ^*^ p<0.05. WT: n=3; *Mcl1*^Δhep^ : n=4. C) Visualization of eccDNA isolated from *Mcl1*^Δhep^ liver by electron microscopy. Scale bar: 100nm. D) Immune cells isolated from *Mcl1*^Δhep^ liver showing higher p-p65 and STING levels. E) Densitometric analysis of Western blot for p-p65 level normalized to CD45 level from 3 independent experiments. Student’s t-test ^**^ p<0.01. F) Visualization of eccDNA isolated from AML12 cells treated with 0.5μM of staurosporine for 24 hours or 0.5μM of reversine for 48 hours. Scale bar: 250nm & 200nm. G) Induction of *Ifnb1* expression analyzed by qPCR after poly(dGdC) treatment on WT BMDM showed a dose-dependent pattern. Student’s t-test ^*^ p<0.05. n=3. H) BMDMs from WT, *Sting1*^-/-^ and *Cgas*^-/-^ mice were transfected with 10ng/ml of eccDNA, linear genomic DNA, and poly(dGdC). The *Ifnb1* expression was analyzed 12 hours after the transfection. n=3. One-Way ANOVA: ^***^ p<0.001, ^****^ p<0.0001.

EccDNA from HeLa cells has been shown to be a strong immunostimulant (18). Therefore, we tested the effect of eccDNA from hepatocytes on immune cells and compared them with linear DNA and poly(dGdC). To ensure an adequate amount of eccDNA was generated in hepatocytes, we treated AML12 cells (a murine hepatocyte cell line) with staurosporine, using HeLa cells in parallel as a control. Treatment with staurosporine for 24 hours strongly increased the percentage of apoptotic cells in AML12 and HeLa cells (Fig. S3B&C). EccDNA from these cells were isolated and visualized by electron microscopy (Fig. 2F; Fig. S3D). Similarly, treatment of AML12 cells over 48 hours with reversine, a reagent known to induce lagging chromosomes and micronuclei formation, also resulted in eccDNA generation (Fig. 2F). Of note, reversine treatment did not induce extensive apoptosis in AML12 cells as in staurosporine treatment. To compare the immune-stimulating potency of eccDNA with linear DNA on immune cells, we generated bone marrow-derived macrophages (BMDMs) from 2-month-old mice of different genotypes (Fig. S3E&F). Poly(dGdC) was able to induce *Ifnb1* expression in WT BMDMs in a dose-dependent manner (Fig. 2G). At a dosage in which both poly(dGdC) and linear DNA were unable to induce *Ifnb1* expression in WT BMDMs, eccDNA from AML12 cells induced significantly higher expression of *Ifnb1* and *Tnfa* (Fig. 2H, Fig. S3G). A similar result was observed with HeLa cell eccDNA which induced a much stronger expression of *Ifnb1* and *Tnfa* in WT BMDM (Fig. S3H). Such induction was strongly dependent on the cGAS-STING pathway, as a deletion of either cGAS or STING abolished *Ifnb1* and *Tnfa* inductions by eccDNA (Fig. 2H & Fig. S3G&H). These results confirmed that eccDNA had stronger immunostimulating properties and their activities were dependent on both cGAS and STING.

### CirSeq indicates that *Mcl1*^Δhep^ livers have a higher quantity of eccDNA which originates from all chromosomes

To study further the characteristics of eccDNA from *Mcl1*^Δhep^ livers, we performed circular sequencing (cirSeq) using an established protocol (22). In contrast to the silica-column-based extraction method, this protocol enabled the detection of both large and small circular DNAs (17, 22, 23). Rolling circle amplification was performed on the purified circular DNA followed by 150bp pair-ended illumina sequencing (Fig. 3A). The average sequencing depth per sample was 10.01 million reads and 99.76% of the reads were mapped to the reference genome. Via the identification of the regions with high abundance of mapped reads, together with the presence of breakpoints, split reads and outward-facing discordant read pairs, circular DNA structures could be identified. We were able to identify eccDNA of various sizes originating from different regions of the genome. For example, a 1,116bp eccDNA was found to originate from chr1:57,239,706 - 57,240,822 within the long non-coding RNA BC055402 locus of a *Mcl1*^Δhep^ liver sample (Fig. 3B). We detected an average of 1,353 unique eccDNA in WT livers, but more than 10 times the amount in *Mcl1*^Δhep^ livers, with an average of 14,835 eccDNA per sample (Fig. 3C). This result was consistent with the observation using a different method reported above (Fig. 2B). Despite an increase in the number of eccDNA in *Mcl1*^Δhep^ samples, the average length of circular DNA was significantly shorter (Fig. 3D). We observed that the majority of the eccDNA had a length between 400bp and 6000bp, which accounted for 85.18% of eccDNA in the WT and 95.81% in the *Mcl1*^Δhep^ group, respectively (Fig. 3E). By aligning the sequences to each chromosome, we detected that eccDNA originated from all chromosomes, with the abundance proportional to the size of each chromosome (Fig. 3F). Of note, there was no significant difference in the length of eccDNA originating from different chromosomes. Looking into the content of eccDNA from WT and *Mcl1*^Δhep^ livers, over 99.6% of the eccDNA contained intergenic sequences or partial genes (Fig. 3G). EccDNA carrying partial genes from WT and *Mcl1*^Δhep^ samples originated mostly from intronic regions, though the level was slightly higher in the latter case (65.37% vs. 70.95%). In WT samples, 17 out of 5,413 detected eccDNA carried full genes (0.31%), while in *Mcl1*^Δhep^ samples, 44 out of 58,860 eccDNA carried full genes (0.075%) (Table S1). These results suggested that the increased eccDNA in *Mcl1*^Δhep^ livers was mostly composed of smaller eccDNA which originated from all chromosomes.

**Fig. 3.**
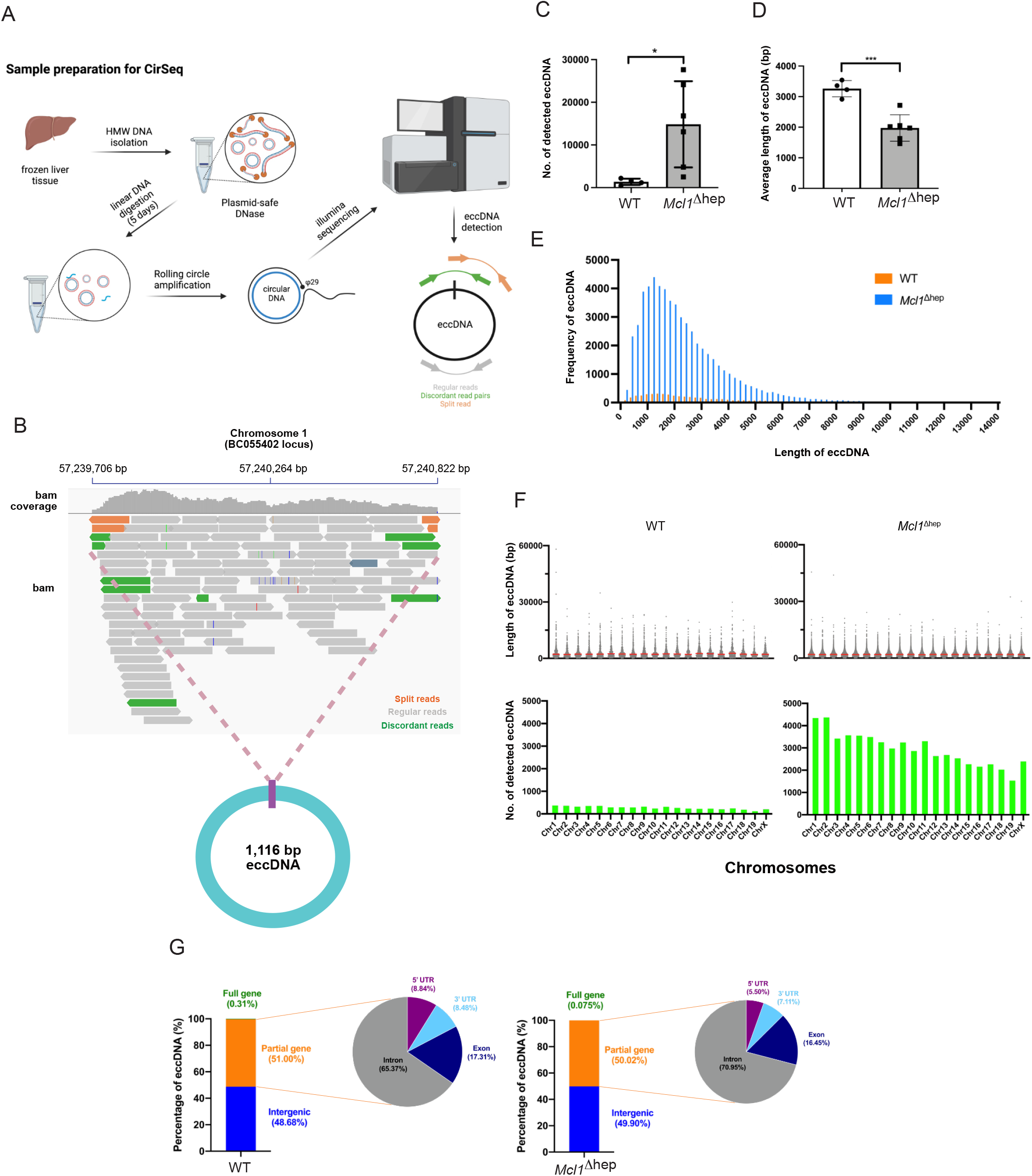
CirSeq shows that eccDNA from *Mcl1*^Δhep^ liver is shorter and comes from all chromosomes. A) Schematic representation showing the steps of sample preparation and CirSeq. B) Example of a 1,116 bp eccDNA identified on chromosome 1 at the BC055402 locus of a *Mcl1*^Δhep^ liver sample. C) Number of eccDNA detected in each liver sample. D) Average length of eccDNA (bp) detected from each liver sample. WT: n=4; *Mcl1*^Δhep^ : n=6. Student’s t-test: ^*^ p<0.05, ^**^ p<0.01. E) Plotting of frequency of eccDNA according to different size (bp). Bin size: 200 bp. n=4. F) Size and number of detected eccDNA from each chromosome. Upper: Each dot represents one detected eccDNA. The red line indicates the mean. Lower: Data were summarized from 4 samples from each group. G) The content of detected eccDNA was analyzed and the percentages of full gene, partial gene or intergenic containing eccDNA were shown.

### Micronuclei contain more abundant eccDNA

DNA damage in micronuclei has been associated with chromothripsis, particularly in cells re-integrating the DNA fragments in micronuclei back into the genome (24). During chromothripsis, cells acquire massive genomic rearrangements through DNA fragmentation and rejoining of DNA in a random manner (25). Since we observed an increase in micronucleated hepatocytes in *Mcl1*^Δhep^ mice (Fig. 1D), we hypothesized the presence of these structures is associated with the increase in eccDNA. To test this hypothesis, we treated AML12 cells with reversine for 48 hours to induce micronuclei (Fig. S4A&B). Quantification of γH2AX fluorescence intensity showed that micronuclei exhibited more severe DNA damage compared to primary nuclei (Fig. S4C). By labeling the cells with EdU, we observed a portion of micronuclei actively underwent DNA replication (Fig. S4D). To study micronuclei specifically, we performed micronuclei isolation based on a method using sucrose gradient centrifugation to separate micronuclei- and primary nuclei-rich fractions (26). Based on the DNA intensity (Hoechst 33343) and size (forward-scattered light), we sorted primary nuclei (PN) and micronuclei (MN) from both AML12 and HEK293T cells using flow cytometry and performed cirSeq (Fig. S5A). Interestingly, micronuclei from AML12 cells showed much higher eccDNA levels compared to primary nuclei (Fig. 4A). Similarly to the distribution observed in livers of *Mcl1*^Δhep^ mice, the majority of eccDNA was below the length of 6000bp (Fig. 4B). Alignment of the sequences indicated that also these eccDNA originated from all chromosomes (Fig. S5B). A similar pattern was observed when comparing micronuclei and primary nuclei from HEK293T cells (Fig. 4A&B; Fig. S5D). These observations suggest that micronuclei bear more abundant eccDNA irrespective of the cell types. Furthermore, by comparing eccDNA from micronuclei and primary nuclei, we observed that micronuclei tended to promote the generation of small eccDNA (Fig. 4C; Fig. S5C&E). These patterns suggested that the more abundant small eccDNA observed in *Mcl1*^Δhep^ compared to WT livers were due to higher levels of micronucleated hepatocytes. By analyzing the sequences of eccDNA identified in PN and MN, MN appeared to contain slightly more eccDNA with partial genes (Fig. S5F&G). Delving into the partial gene-containing eccDNA, the ones with intronic sequences were more abundant in MN compared to PN (Fig. 4D).

**Fig. 4.**
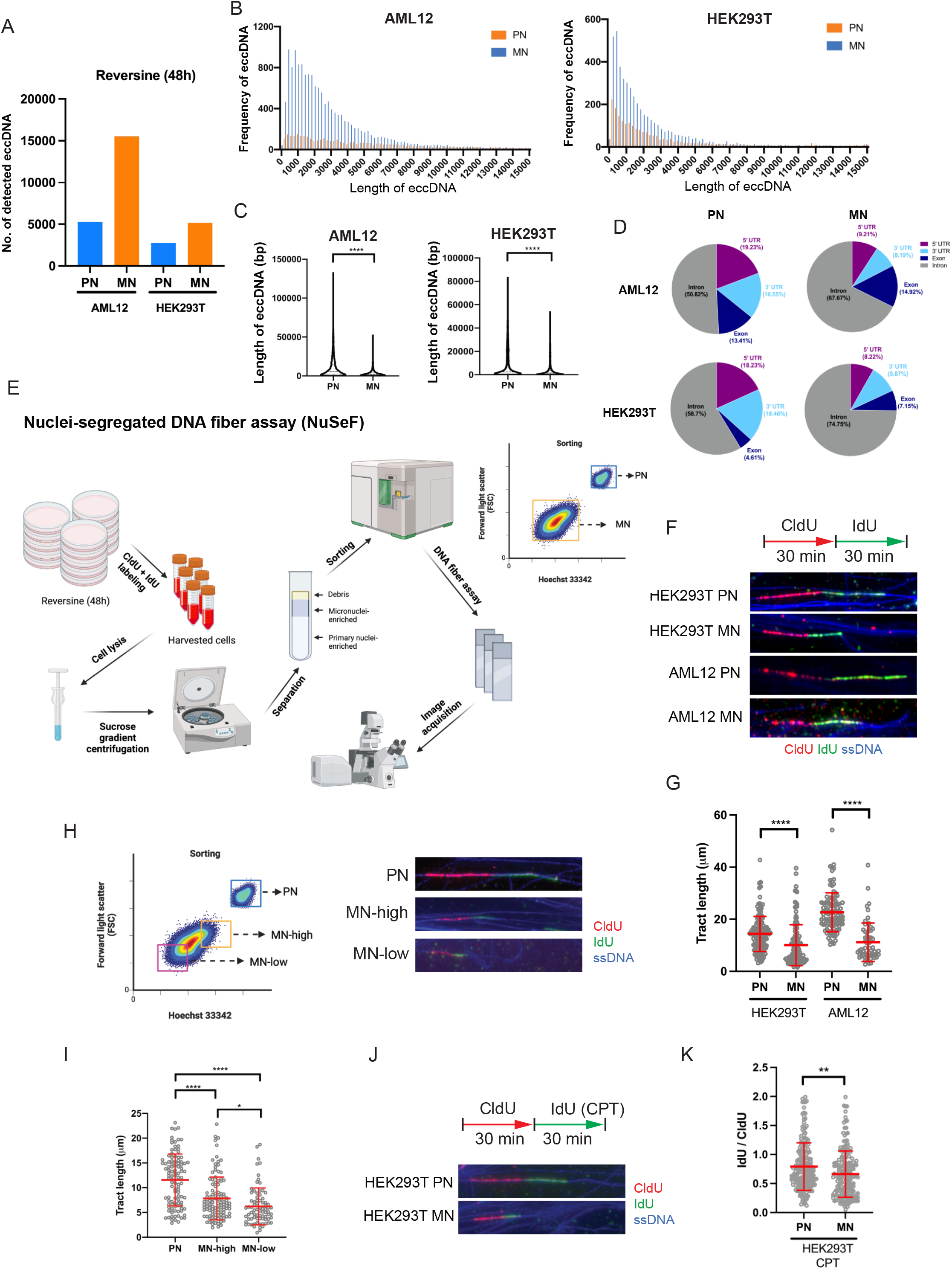
Micronuclei have more abundant eccDNA than primary nuclei. A) CirSeq performed on sorted primary nuclei and micronuclei. The number of detected eccDNA was compared. B) Plotting of frequency of eccDNA detected in AML12 and HEK293T primary nuclei and micronuclei according to different sizes (bp). Bin size: 200bp. C) Comparison of the length of eccDNA (bp) between primary nuclei and micronuclei. D) Comparison of the distribution of partial-gene-containing eccDNA which harbor 5’ UTR, 3’UTR, intronic and exonic sequences. E) Schematic of NuSeF assay. F) NuSeF assay comparing the DNA fibers from primary nuclei and micronuclei. CldU and IdU were pulse-labelled for 30 minutes each. Single-stranded DNA (ssDNA) staining was used to visualize the integrity of DNA. G) Tract length (CldU+IdU) was measured. Each dot represented a single fiber. H) NeSeF assay showing DNA fiber length was decreasing from primary nuclei (PN), large micronuclei (MN-high) and small micronuclei (MN-low) from HEK293T cells. I) Tract length (CldU+IdU) was measured. Each dot represented a single fiber. J) Cells were treated with reversine for 48 hours. NuSeF assay was performed with treatment with/without CPT at the IdU labeling step. K) The ratio of IdU to CldU was measured to study the frequency of fork-stalling events under CPT treatment. Student’s t-test: ^**^ p<0.01, ^***^ p<0.00, ^****^ p<0.0001 (C, I & K). One-Way ANOVA: ^*^ p<0.05, ^****^ p<0.0001 (G).

### Nuclei-segregated DNA fiber (NuSeF) assay reveals the intrinsic defects in micronuclei showing stronger replication stress

To better understand how micronuclei promote eccDNA generation, we developed a new method called NuSeF assay (Fig. 4E). This approach allowed us to distinguish DNA fibers originating from primary nuclei and micronuclei after separation by FACS. Using NuSeF assay, DNA fibers from micronuclei of AML12 and HEK293T cells showed much shorter total tract length (CldU + IdU) (Fig. 4F&G) compared to their corresponding primary nuclei. The reduction was observed in both the CldU and IdU tract lengths, though the IdU to CldU ratio was not altered (Fig. S6A-C). This suggested that micronuclei had higher replication stress which resulted in overall replication fork slowing. Such a pattern was consistent in both cell types. Interestingly, we observed an inverse correlation between the size of nuclei (primary nuclei vs. micronuclei-high vs. micronuclei-low) and the total DNA tract length (Fig. 4H&I). Similarly to primary nuclei, micronuclei also responded to camptothecin (CPT) treatment, showing a reduced total tract length upon induction of replication stress (Fig. S6F). Treatment with CPT also resulted in a significant reduction of the IdU to CldU ratio in micronuclei compared to primary nuclei (Fig. 4J&K). This indicated that micronuclei were more susceptible to replication stress which might explain the observed higher DNA damage in these structures (Fig. S4C). Our results suggested that, although micronuclei still retained the replication properties observed in primary nuclei, intrinsic defects in micronuclei such as their increased susceptibility to replication stress could be a source of increased eccDNA generation.

### EccDNA is increased in mouse models of liver diseases with high tumor incidence

We then aim to discern whether the increase in eccDNA was specific to the *Mcl1*^Δhep^ mouse model. We performed cirSeq in several mouse models of liver diseases including a model of polycystic liver disease (JNK1/2^Dhep^), a model of fatty-liver disease (fast-food diet), and a model of oncogene activation (c-MYC overexpression) (Fig. 5A) (27-29). JNK1/2^Dhep^ mice at 12 months displayed multiple cystic structures in the liver. However, this phenotype was not associated with an increase in eccDNA (Fig. 5B). Mice fed with a fast-food diet developed macrovesicular steatosis but did not show any changes in eccDNA levels. In contrast, high eccDNA levels were observed only in *Mcl1*^Δhep^ and *Myc*^OE^ livers. These models had reduced average size of eccDNA, suggesting that the increased eccDNA was mostly small eccDNA. Interestingly, both *Mcl1*^Δhep^ and *Myc*^OE^ mouse lines had much higher tumor incidence (∼50% in 12-month-old *Mcl1*^Δhep^ mice and 100% in *Myc*^OE^ mice) compared to other models (0% in JNK1/2^Dhep^ mice and 2.5% in 12-month-old fast-food diet mice) (27-29). These data suggested a correlation between high eccDNA levels and HCC incidence.

**Fig. 5.**
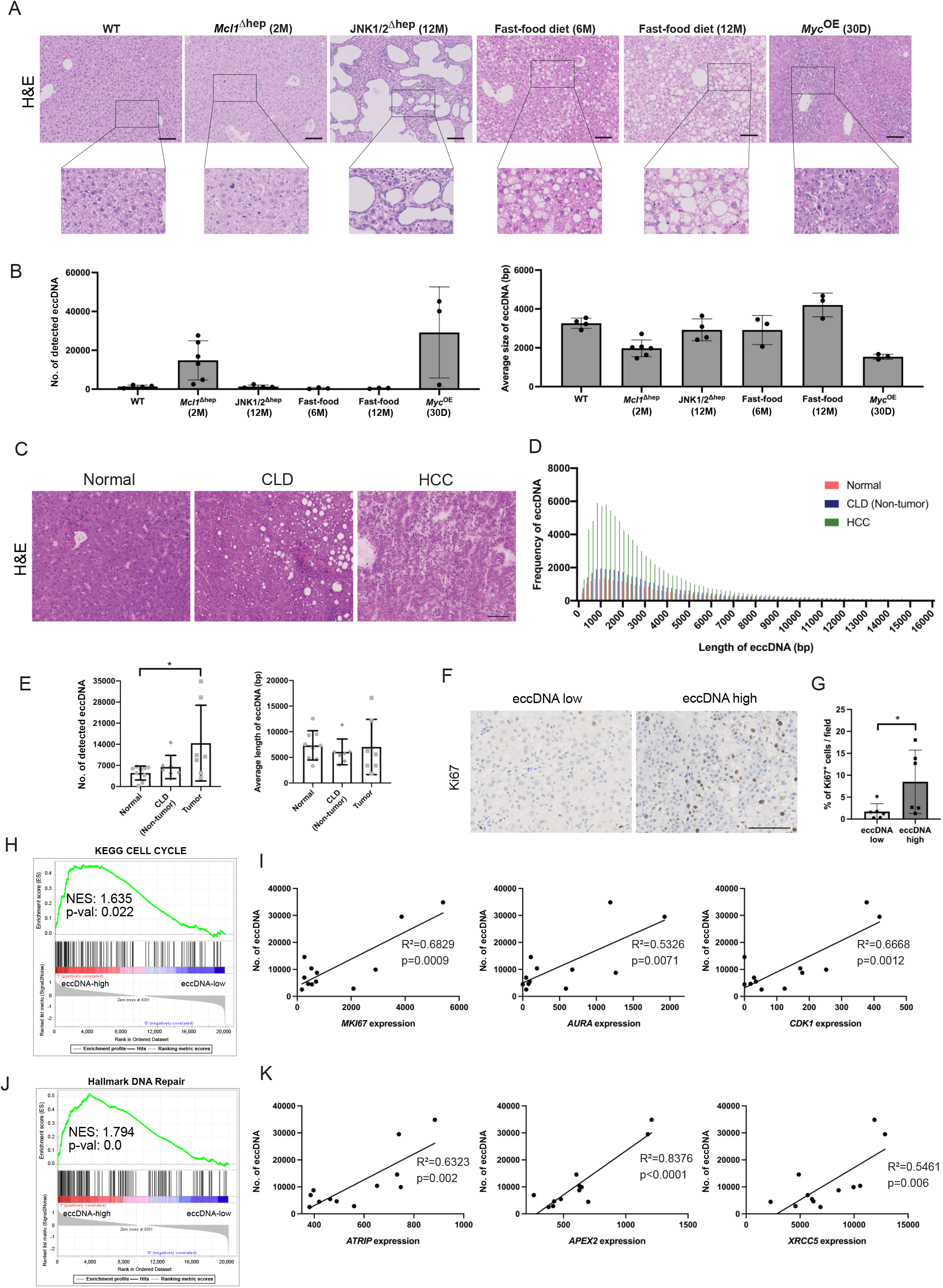
eccDNA levels are increased in mouse models with high tumor incidence and HCC patient samples. A) Histology of mouse models of various liver diseases: 2-month-old *Mcl1*^Δhep^ liver (chronic liver injury); 12-month-old JNK1/2^Δhep^ mice (polycystic liver disease); 30-day-old LAP-tTa/tetO-c-MYC mice (MYC oncogene activation); 6-month and 12-month fast-food diet treated mice (fatty liver disease). Scale bar: 100μm. B) CirSeq was performed from the liver tissues of these mice. The average number of detected eccDNA and their average size were compared. C) Histology of patient tissues from normal, CLD (non-tumor area) and HCC livers. Scale bar: 100μm. D) Plotting of frequency of eccDNA according to different size (bp). Bin size: 200bp. n=7. E) Total number and average length of detected eccDNA. Each dot represented one sample. Student’s t-test: ^*^ p<0.05. F) Staining of Ki67 on HCC samples from the eccDNA-high and eccDNA-low groups. Scale bar: 100μm. G) Quantification of the percentage of Ki67^+^ cells per field on the eccDNA-high and eccDNA-low CLD and HCC samples. Student’s t-test: ^*^ p<0.05. H) GSEA using RNAseq data comparing the eccDNA-high and eccDNA-low groups showing an enriched ‘Cell Cycle’ signature in the eccDNA-high group. I) Correlation of eccDNA levels from CLD and HCC samples to their corresponding expression of proliferation-associated genes (*MKI6, AURA* and *CDK1*) using normalized counts. J) GSEA showing an enriched ‘DNA Repair’ signature in the eccDNA-high group. K) Correlation of eccDNA levels to their corresponding expression of DNA repair-related genes (*ATRIP, APEX2* and *XRCC5*) using normalized counts. Pearson’s correlation coefficient and R^2^ were calculated in all the correlation tests.

### EccDNA levels are increased in human CLD and HCC tissues

To study if elevated eccDNA levels might be relevant for tumorigenesis in humans, we examined eccDNA levels in HCC and the adjacent CLD tissues in comparison to non-diseased liver tissues (Fig. 5C). CirSeq revealed that the eccDNA levels were already increased in CLD, and significantly elevated in HCC tissues (Fig. 5D&E). We observed that the majority of eccDNA had a size below 8,000bp. The largest detected circular DNA in patient samples was 724,600 bp which was still much smaller than the expected size observed in extrachromosomal DNA (ecDNA), a class of circular DNA frequently detectable in tumor cells (23). To determine whether some of these circular DNA would carry complete genes, the cirSeq sequences were aligned to the human genome (GRCh38), where we were able to identify eccDNA carrying at least one complete gene in all samples, with an average of 95, 62, and 73 eccDNA per sample in the normal, CLD and HCC group respectively. To analyze if there was a correlation between the presence of full genes in eccDNA to their expression, we performed total RNAseq. A comparison of samples that contained full genes present in eccDNA with the rest of the samples did not reveal a significant upregulation of these genes, suggesting that these genes were most likely not being amplified in the circular form (Fig. S7A-C). Since eccDNA levels were increased in CLD and HCC tissue samples, we divided these samples into eccDNA-high and eccDNA-low groups based on their eccDNA levels. We observed that the percentage of Ki67^+^ cells in the eccDNA-high group was significantly higher than in the eccDNA-low group (Fig. 5F&G). EccDNA-high samples also had an increased tendency of CD11b^+^ cell infiltration (Fig. S8A). We then performed GSEA and observed a significant enrichment of the ‘Cell Cycle’ signature in the eccDNA-high group (Fig. 5H). By performing correlation analyses, we found that the level of eccDNA was strongly correlated to the expression of proliferation-associated genes, e.g. *MKI67* (gene encoding Ki67), *AURKA* (gene encoding aurora kinase A), and *CDK1* (gene encoding CDK1) (Fig. 5I). In line, a statistically significant correlation was found between the eccDNA levels and the percentage of Ki67^+^ cells (Fig. S8B). Furthermore, we observed an enrichment of the ‘Hallmark DNA Repair’ signature in the eccDNA-high group (Fig. 5J). A strong correlation was also observed between eccDNA levels and the expression of different genes in several DNA repair mechanisms: *ATRIP* which is essential for recruiting ATR in response to double-strand break as part of the DNA damage response (DDR); *APEX2* which encodes an apurinic/apyrimidinic endodeoxyribonuclease that functions in the DNA base excision repair; XRCC5 which encodes Ku80 that together with Ku70 binds to DNA double-strand breaks and promotes non-homologous end-joining (NHEJ) (Fig. 5K; Fig. S8C). These results suggested that patient samples with strong proliferation, and consequent increased replication stress and DNA damage, tended to generate higher levels of eccDNA.

Deletion of *Sting1* reduces inflammation and tumor incidence in *Mcl1*^Δhep^ mice Since we observed that the effect of eccDNA on immune cells in inducing *Ifnb1* and *Tnfa* expression was dependent on the cGAS-STING pathway (Fig. 2H & Fig. S3H), we then sought to determine if the phenotype of *Mcl1*^Δhep^ mice would be affected by blocking this pathway. To achieve this, we crossed *Sting1*^-/-^ mice with *Mcl1*^Δhep^ mice to generate a *Mcl1*^Δhep^ *Sting1*^-/-^ mouse line. We confirmed the knockout status of the *Sting1*^-/-^ mice by Western blot and observed that splenocytes from these mice were not able to induce IRF3 phosphorylation after DMXAA (murine STING1 agonist) stimulation (Fig. S9A). Interestingly, *Mcl1*^Δhep^ *Sting1*^-/-^ mice showed no significant difference in liver damage (serum ALT), level of apoptosis (cl. Caspase 3), proliferation (Ki67), or DNA damage (γH2AX) when compared to *Mcl1*^Δhep^ mice (Fig. 6A&B). Given the expression pattern of STING in the liver (Fig. S2C), the effect of inhibiting the STING pathway was assumed to affect immune cells. To test this, total RNAseq and gene signature analyses were performed on *Mcl1*^Δhep^ *Sting1*^-/-^ liver samples using *Mcl1*^Δhep^ mice as a control. Interestingly, over-representation analysis (ORA) indicated that the majority of downregulated signatures in the *Mcl1*^Δhep^ *Sting1*^-/-^ liver were related to immune repones (Fig. 6C). Various ISGs were found to be significantly downregulated upon *Sting1* deletion (Fig. S9B). Using GSEA, we observed a depletion of signatures involved in immune cell chemotaxis in *Mcl1*^Δhep^ *Sting1*^-/-^ mice e.g. macrophage migration and granulocyte chemotaxis (Fig. S9C). In line, quantification of F4/80-positive cells showed that STING knockout reduced the number of macrophages/Kuffer cells in the *Mcl1*^Δhep^ liver (Fig. 6B). Since we observed a reduced immune response upon *Sting1* deletion, we compared cohorts of these mice at the age of 12 months. Interestingly, *Mcl1*^Δhep^ *Sting1*^-/-^ mice showed a reduced tumor incidence of 33.33% compared to 56.52% in the *Mcl1*^Δhep^ group (Fig. 6D&E). These results suggested that deletion of STING reduced immune response and that such a disrupted inflammatory microenvironment led to a reduction in the tumor incidence, highlighting the importance of the cGAS-STING pathway in the eccDNA-mediated inflammation in *Mcl1*^Δhep^ livers.

**Fig. 6.**
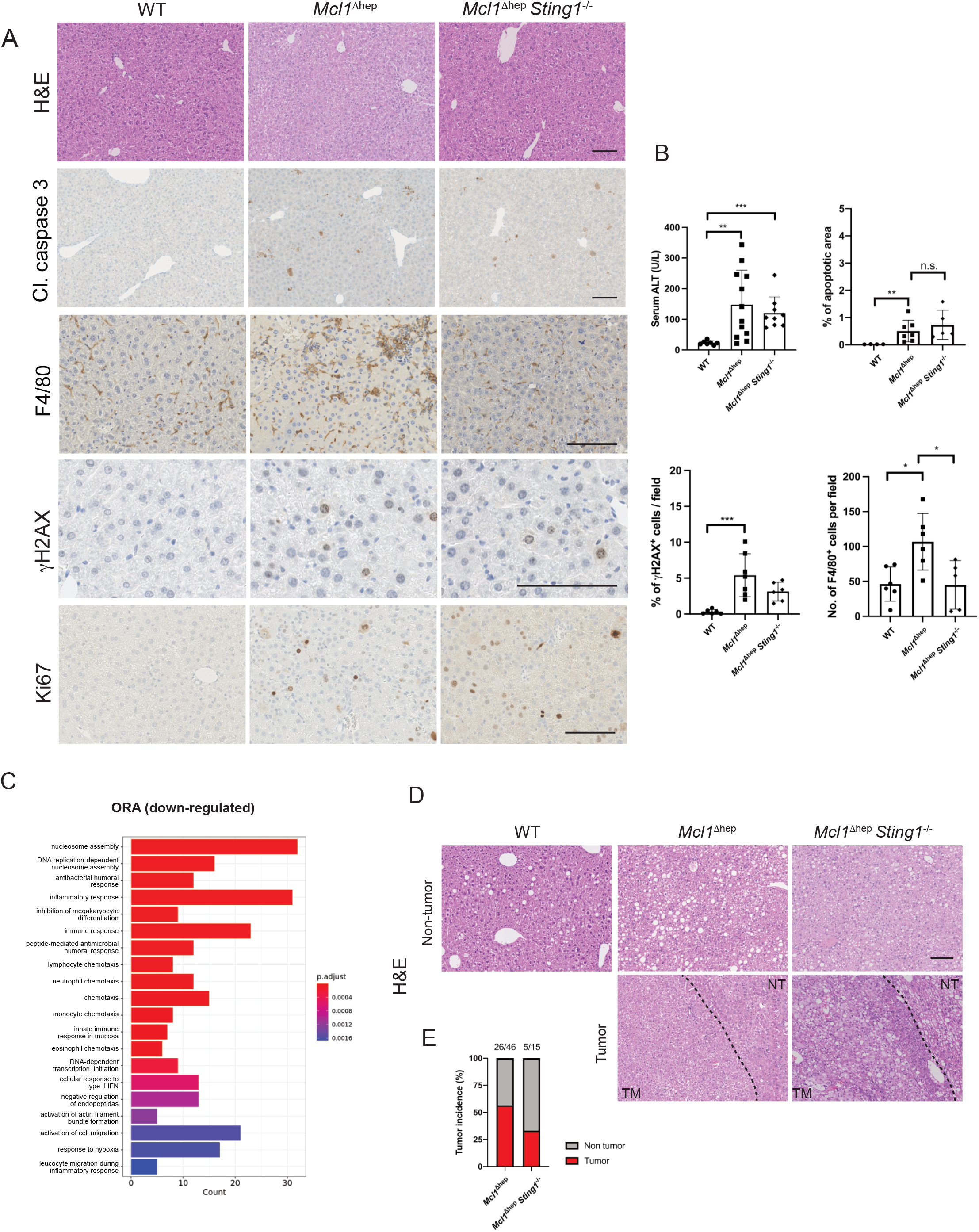
STING knockout reduces immune cell infiltration and tumor incidence in *Mcl1*^Δhep^ mice. A) Histological analyses of 2-month-old mice showed no differences in apoptosis (cleaved caspase 3), DNA damage (γH2AX) and proliferation (Ki67) but a reduction in macrophage infiltration (F4/80) in *Mcl1*^Δhep^ *Sting1*^-/-^ mice. B) Measurement of serum ALT levels from blood samples collected at the 2-month time point. Quantification of the cleaved caspase 3 staining as a percentage of total area. Quantification of the number of γH2AX^+^ hepatocytes per field. Quantification of the number of F4/80^+^ cells per field. Student’s t-test: ^*^ p<0.05, ^**^ p<0.01, ^***^ p<0.001. C) *Mcl1*^Δhep^ *Sting1*^-/-^ mice showed significant downregulation of gene signatures involved in immune responses and immune cell recruitment. ORA: over-representation analysis. D) Histology of 12-month livers showing the tumor (TM) area and non-tumor area (NT). Scale bar: 100μm. E) Tumor incidence showing the percentage of mice in the corresponding genotypes that displayed at least one tumor in the liver.

## Discussion

It has been two decades since the presence of micronuclei in human CLD and HCC tissues was described (30). However, it remains unclear how the presence of micronuclei is connected to HCC carcinogenesis. The results of this study shed new light on this old observation and suggest a strong connection with eccDNA generation, immune response, and liver carcinogenesis. Micronuclei can arise from chromosome mis-segregation, but in cells experiencing replication stress, micronuclei can also contain chromatin fragments caused by DNA damage (10). Using *Mcl1*^Δhep^ mice as a CLD model, DNA fiber assay indicated increased replication stress in MCL1-deficient hepatocytes. Prolonged replication fork stalling often causes fork collapses leading to DNA double-stranded breaks (31). The high replication stress from the apoptosis-triggered hyperproliferation in MCL1-deficient hepatocytes can in turn generate higher DNA damage and micronuclei which will eventually promote apoptosis in a positive feedback loop manner. Although apoptosis has been shown to promote eccDNA generation, other studies also show their biogenesis in relation to other stress conditions or DNA damage (18, 19). Our data suggested that eccDNA generation and accumulation occurred already before apoptosis, through micronuclei formation.

To gain insight into the relationship between micronuclei and eccDNA formation, we developed a method that allows us to study replication stress directly in micronuclei. This new approach, designated NuSeF assay, allowed us to distinguish DNA fibers originating from micronuclei and primary nuclei, in contrast to the conventional protocol which examines the bulk DNA fibers. NuSeF assay revealed that micronuclei were more susceptible to replication stress, as indicated by replication fork slowing, thus corroborating overall higher DNA damage levels in micronuclei compared to primary nuclei discovered by imaging methods. Using cirSeq, eccDNA was detected in both primary nuclei and micronuclei. The fact that eccDNA was detected in primary nuclei may explain the previous report that eccDNA was found in various healthy tissues (17). Presumably, the main source of eccDNA from healthy tissue is from primary nuclei, as healthy cells generally do not contain micronuclei. Though micronuclei contained more abundant eccDNA, as shown in this study, their eccDNA is much smaller. This can partly reflect the more abundant but smaller eccDNA observed in *Mcl1*^Δhep^ livers, due to the increased proportion of micronucleated cells. Importantly, the micronucleated hepatocytes in *Mcl1*^Δhep^ livers and the micronuclei-containing AML12 cells after reversine treatment were non-apoptotic. One limitation of our result was that we could not completely exclude a contribution of apoptosis in eccDNA biogenesis in the *Mcl1*^Δhep^ model since apoptosis was one of the effects of MCL1 deletion in hepatocytes. As discussed above, excessive DNA damage can eventually promote apoptosis, and as such, the more likely scenario is that eccDNA are generated after micronuclei formation and the generation is further enhanced by apoptosis later. This is in line with the current understanding that eccDNA can be produced by multiple mechanisms (32). Further application of the NuSeF assay, in combination with cirSeq, on cells exposed to various genotoxic insults or in experimental designs to restore replication defects within micronuclei, will help a better understanding of the micronuclei biology and may reveal additional mechanisms of eccDNA generation.

The circular nature of eccDNA is likely to provide additional stability and therefore increase the potency in inducing immune response (18). We observed that even at low dosages, eccDNA can induce *Ifnb1* expression in BMDMs. The potency can also be an effect of eccDNA resistance to intracellular exonuclease such as TREX1, which has an inhibitory function for the activation of the cGAS-STING pathway (33). However, it is still unclear whether eccDNA is actively secreted by damaged hepatocytes, or if they are released during cell death. The ability of eccDNA to induce a strong response in immune cells suggests that they are acting as DAMP molecules. Furthermore, it is also unclear if eccDNA is secreted locally to activated adjacent immune cells, or if they are in the circulation as part of the circulating DNA. In fact, cell-free eccDNA has been reported in the plasma of both mice and humans which suggests a possibility to reach distant targets (34). Further studies are required to elucidate if they are involved in a systemic immune response.

Although an increase in DNA damage may eventually increase the number of accumulated mutations which ultimately can drive neoplastic transformation, we observed in this study an essential role of the immune cells in the carcinogenesis process in the *Mcl1*^Δhep^ model. Our results suggested a crosstalk between hepatocytes and the microenvironment, with eccDNA serving as a link connecting the increased DNA damage in hepatocytes to the activation of the cGAS-STING pathway in the immune cells. This connection is essential since hepatocytes do not express STING (12, 14). More importantly, inhibition of this pathway reduced the overall immune cell infiltration in *Mcl1*^Δhep^ livers and reduced the tumor incidence at 12 months. Inflammation has been long known to be associated with HCC development (35). Therefore, our results suggest that carcinogenesis in *Mcl1*^Δhep^ mice, which very faithfully recapitulate the CLD pathophysiology, is mediated through a non-cell-autonomous mechanism.

In the current study, we showed that the effect of eccDNA on immune cells was dependent on both cGAS and STING. In animal models receiving HFD (high-fat diet) or MCD (methionine- and choline-deficient diet), constitutive knockout of STING or selective knockout of STING in myeloid cells both led to reduced NASH phenotype, which includes attenuated liver steatosis, inflammation, and fibrosis (12, 14). These studies also highlighted the observation of a lack of STING expression in hepatocytes, as confirmed in the current study. Such observation also supports the concept that hepatocytes are susceptible to HBV infection due to a lack of the DNA sensing pathway and their inability to induce type 1 interferon response after viral infection (13). Therefore, the postulated role of STING in eliciting a senescence-associated secretory phenotype (SASP) might be important in other cell types, but may not be relevant in hepatocytes (36). Due to the different spatial expressions of cGAS and STING, selective inhibition of these components may result in different outcomes. Several studies reported a non-canonical role of cGAS in regulating senescence, cell death, and DNA repair (37-40). Similarly, non-canonical activation of STING after DNA damage has been reported through the recruitment of ATM and IFI16 leading to the activation of the downstream NF-κB signaling (41). The relevance of these non-canonical functions in hepatocytes especially in the pathogenesis of liver diseases still requires further elucidation.

This study provided evidence that eccDNA, generated due to an accumulation of micronuclei, act as a link to promote crosstalk between hepatocytes and immune cells to maintain immune response in the liver. Our results showed that carcinogenesis in *Mcl1*^Δhep^ mice was mediated by a non-cell-autonomous mechanism, with the crosstalk between hepatocytes and immune cells playing a pivotal role. NuSeF assay is a novel approach to enable the study of replication fork dynamics in micronuclei leading to the discovery of higher replication stress in micronuclei.

## Materials and Methods

### Mouse lines and patient materials

*Mcl1*^Δhep^ mice were generated by crossing Alb-Cre transgenic mice with *Mcl1* floxed mice as described previously (6). Liver of JNK1/2^Δhep^ mice, mice treated for 6 or 12 months of fast-food diet, and *Myc*^OE^ mice were obtained as described previously (27-29). *Sting1*^-/-^ mice were kindly provided by Prof. Dr. Thorsten Buch (LASC, UZH). *Mcl1*^Δhep^ *Sting1*^-/-^ mice were generated by crossing *Mcl1*^Δhep^ mice with *Sting1*^-/-^ mice. *Cgas*-/-mice were purchased from JAX. WT, *Mcl1*^Δhep^ and *Mcl1*^Δhep^ *Sting1*^-/-^ mice were analyzed at the age of 2 and 12 months. Human liver samples, snap-frozen or formalin-fixed, paraffin-embedded (FFPE), were provided by the bio-bank of the Department of Pathology and Molecular Pathology, University Hospital Zurich, for morphological and molecular analyses.

### Nuclei-segregated DNA fiber (NuSeF) assay

Cells were pulse-labeled with two thymidine analogs, chlorodeoxyuridine (CldU) and iododeoxyuridine (IdU) for 30 minutes each. Labeled cells were harvested and treated with cytochalasin B for 30 minutes on ice. Primary nuclei and micronuclei were enriched in fractions by sucrose gradient centrifugation. FACS was performed on these enriched fractions to obtain pure primary nuclei and micronuclei. Purified nuclei were lysed and the DNA content was spread on glass slides. The slides were then incubated with anti-CldU (BD Bioscience #347580) and anti-IdU (abcam #ab6326) primary antibodies, followed by secondary antibodies. Single-stranded DNA (ssDNA) (DSHB TROMA-III) counterstain was performed to visualize the integrity of DNA fibers. The fiber tract length was quantified and compared.

### Primary hepatocyte and immune cell isolation from the liver

Hepatocyte isolation was performed according to the proposed protocol (42). Briefly, mice were first anesthetized by ketamine (100mg/kg) and xylazine (16mg/kg). The liver was first perfused with the pre-perfusion buffer (0.5mM EDTA/20mM HEPES/HBSS) for 5 minutes and then switched to perfusion solution (20mM HEPES / 1X Penicillin/Streptomycin / 3mM CaCl_2_ / DMEM/F12 / 0.2mg/ml Liberase (Roche #05401127001)) for 10 minutes. The liver was then transferred to a petri dish with 10ml of Wash solution (4% FBS / 1X Penicillin/Streptomycin / Willian’s E medium) and hepatocytes were gently released from the liver using forceps. The Wash solution containing cells was filtered through a 70μm cell strainer (Falcon #352350) and stored on ice. The cell suspension was centrifuged at 20g for 3 minutes. The pellet was used for hepatocyte isolation and the supernatant, which contained non-parenchymal cells, was transferred to a new 50ml Falcon tube for immune cell isolation. For hepatocyte purification, the pellet was resuspended with 10ml of Wash solution and mixed with 10ml of 90% percoll solution (Cytiva #GE-17-0891-01) and centrifuged at 200g for 10 minutes. This step was repeated one more time to increase the purity of viable cells. The pellet was then resuspended in Wash solution and seeded at a density of 3×10^5^ cells/well in a collagen pre-coated 24-well plate. For isolating immune cells, the volume was brought to 50ml using the Wash solution. The solution was centrifuged at 20g for 3 minutes and the supernatant was transferred to a new 50ml Falcon tube. This step was repeated for additional two times. The transferred supernatant was then centrifuged at 2000 rpm for 5 minutes. The immune cell pellet was resuspended in 10ml of 36% Percoll solution and centrifuged at 2000rpm for 20 minutes at 4°C. RBCs were lysed with 1X RBC lysis buffer (G Biosciences #786-649) at room temperature for 5 minutes. The immune cells were washed once with 10ml of PBS and pelleted followed by snap-freezing and storage. The purity of the isolated cells was examined using Western blot for hepatocyte- and immune cell-specific markers (albumin and CD45) respectively.

### DNA fiber assay

For DNA fiber assay on primary hepatocytes, primary hepatocytes were isolated as described above. Hepatocytes were seeded in collagen-coated 6-well plates at a density of 20% after isolation. Hepatocytes were cultured for 24 hours before the assay. On the day of the experiment, hepatocyte medium was removed from the wells and first replaced with 19mM CldU in fresh medium followed by exchanging the medium with 28.24mM IdU in fresh medium at 30-minute intervals. The wells were washed twice with pre-warmed PBS in between CldU and IdU-containing media. After the IdU incubation, cells were washed twice with cold PBS and trypsinized. Cells were harvested and stored on ice until DNA spread. Approximately 1,000 cells in 3μl were transferred onto a glass slide and then mixed with 7μl of lysis buffer (200mM Tri-HCl pH7.4 / 50mM EDTA / 0.5% SDS). After 5 minutes of lysis at room temperature, the slides were tilted at an angle of 45° to promote the spreading of DNA by gravity. The slides were then dried and fixed with methanol/acetic acid (3:1) for 20 minutes at 4°C. Slides were rehydrated in PBS and then denatured with 2.5M HCl for 1 hour at room temperature. These slides were then washed with PBS 5 times and blocked with blocking buffer (2% BSA / 0.1% Tween-20 / PBS) for 1 hour at room temperature. The slides were then incubated with anti-CldU (BD Bioscience #347580) and anti-IdU (abcam #ab6326) primary antibodies overnight at 4°C followed by 2 hours of secondary antibodies at room temperature. Single-stranded DNA (ssDNA) (DSHB TROMA-III) counterstain was performed to visualize the integrity of DNA fibers. The stained slides were mounted with ProLong Gold Antifade mounting medium (Invitrogen #P36930). DNA fiber images were acquired using the Leica DM6 microscope.

## Supporting information

Supplementary figures, tables and detailed methods

## Data availability

Gene expression data are available in the GEO database under accession numbers GSE75730, GSE277012 and GSE277232. All cirSeq raw data are submitted to Sequence Read Archive (SRA) (PRJNA1167186).

### Statistical Analysis

Statistical analyses were performed as indicated in figure legends. For a comparison between the two groups, a Student’s t-test was used. For a comparison of more than two groups, one-way ANOVA was used. For all analyses: ^*^ p<0.05; ^**^ p<0.01; ^***^ p<0.001; ^****^ p<0.0001. Sample numbers (n) were indicated in figure legends. All graphs showed a mean±SD. Each dot in all dot-plots indicated each sample or individual quantified unit. Detailed information was provided in the corresponding figure legend. Statistical tests were performed using GraphPad Prism 10.

Extended technical descriptions are available in Extended Methods.

## Acknowledgments

We would like to thank Prof. Pavel Janscak, Dr. Martin Andrs and Mr. Vinicio Rosano for the technical support of the DNA fiber assays. We would also like to thank Prof. Massimo Lopes, Dr. Jana Krietsch and Mr. Moses Aouami for the preparation of samples for electron microscopy. In addition, we would like to thank Prof. Thorsten Buch and Dr. Filipa Marques Ferreira for providing us with the STING knockout mice.

## Funding

Swiss National Science Foundation (SNSF) project fund Hartmann Müller Foundation

### Author contributions

Conceptualization: LKC, AW Methodology: LKC, JS, NW, ERF, AGH

Investigation: LKC, JS, MEH, RP, ND, GS, PL Visualization: LKC,

Supervision: AW, Writing—original draft: LKC

Writing—review & editing: LKC, AW, PL

## References

1. Rumgay, H., Arnold, M., Ferlay, J., Lesi, O., Cabasag, C.J., Vignat, J., Laversanne, M., McGlynn, K.A., Soerjomataram, I. Global burden of primary liver cancer in 2020 and predictions to 2040. J Hepatol. 77, 1598–606 (2022).

2. Yang, J.D., Hainaut, P., Gores, G.J., Amadou, A., Plymoth, A., Roberts, L.R. A global view of hepatocellular carcinoma: trends, risk, prevention and management. Nat Rev Gastroenterol Hepatol. 16, 589–604 (2019).

3. Ringelhan, M., Pfister, D., O’Connor, T., Pikarsky, E., Heikenwalder, M. The immunology of hepatocellular carcinoma. Nat Immunol. 19, 222–32 (2018).

4. Weber, A., Boger, R., Vick, B., Urbanik, T., Haybaeck, J., Zoller, S., Teufel, A., Krammer, P.H., Opferman, J.T., Galle, P.R., Schuchmann, M., Heikenwalder, M., Schulze-Bergkamen, H. Hepatocyte-specific deletion of the antiapoptotic protein myeloid cell leukemia-1 triggers proliferation and hepatocarcinogenesis in mice. Hepatology. 51, 1226–36 (2010).

5. Nakagawa, H., Umemura, A., Taniguchi, K., Font-Burgada, J., Dhar, D., Ogata, H., Zhong, Z., Valasek, M.A., Seki, E., Hidalgo, J., Koike, K., Kaufman, R.J., Karin, M. ER stress cooperates with hypernutrition to trigger TNF-dependent spontaneous HCC development. Cancer Cell. 26, 331–43 (2014).

6. Boege, Y., Malehmir, M., Healy, M.E., Bettermann, K., Lorentzen, A., Vucur, M., Ahuja, A.K., Bohm, F., Mertens, J.C., Shimizu, Y., Frick, L., Remouchamps, C., Mutreja, K., Kahne, T., Sundaravinayagam, D., Wolf, M.J., Rehrauer, H., Koppe, C., Speicher, T., Padrissa-Altes, S., Maire, R., Schattenberg, J.M., Jeong, J.S., Liu, L., Zwirner, S., Boger, R., Huser, N., Davis, R.J., Mullhaupt, B., Moch, H., Schulze-Bergkamen, H., Clavien, P.A., Werner, S., Borsig, L., Luther, S.A., Jost, P.J., Weinlich, R., Unger, K., Behrens, A., Hillert, L., Dillon, C., Di Virgilio, M., Wallach, D., Dejardin, E., Zender, L., Naumann, M., Walczak, H., Green, D.R., Lopes, M., Lavrik, I., Luedde, T., Heikenwalder, M., Weber, A. A Dual Role of Caspase-8 in Triggering and Sensing Proliferation-Associated DNA Damage, a Key Determinant of Liver Cancer Development. Cancer Cell. 32, 342–59 e10 (2017).

7. Thomas, L.W., Lam, C., Edwards, S.W. Mcl-1; the molecular regulation of protein function. FEBS Lett. 584, 2981–9 (2010).

8. Jackson D, P.A. Replicon Clusters Are Stable Units of Chromosome Structure: Evidence That Nuclear Organization Contributes to the Efficient Activation and Propagation of S Phase in Human Cells. J Cell Biol. 140, 1285–95 (1998).

9. Merrick, C.J., Jackson, D., Diffley, J.F. Visualization of altered replication dynamics after DNA damage in human cells. J Biol Chem. 279, 20067–75 (2004).

10. Di Bona, M., Bakhoum, S.F. Micronuclei and Cancer. Cancer Discov. 14, 214–26 (2024).

11. Kwon, J., Bakhoum, S.F. The Cytosolic DNA-Sensing cGAS-STING Pathway in Cancer. Cancer Discov. 10, 26–39 (2020).

12. Luo, X., Li, H., Ma, L., Zhou, J., Guo, X., Woo, S.L., Pei, Y., Knight, L.R., Deveau, M., Chen, Y., Qian, X., Xiao, X., Li, Q., Chen, X., Huo, Y., McDaniel, K., Francis, H., Glaser, S., Meng, F., Alpini, G., Wu, C. Expression of STING Is Increased in Liver Tissues From Patients With NAFLD and Promotes Macrophage-Mediated Hepatic Inflammation and Fibrosis in Mice. Gastroenterology. 155, 1971–84 e4 (2018).

13. Thomsen, M.K., Nandakumar, R., Stadler, D., Malo, A., Valls, R.M., Wang, F., Reinert, L.S., Dagnaes-Hansen, F., Hollensen, A.K., Mikkelsen, J.G., Protzer, U., Paludan, S.R. Lack of immunological DNA sensing in hepatocytes facilitates hepatitis B virus infection. Hepatology. 64, 746–59 (2016).

14. Yu, Y., Liu, Y., An, W., Song, J., Zhang, Y., Zhao, X. STING-mediated inflammation in Kupffer cells contributes to progression of nonalcoholic steatohepatitis. J Clin Invest. 129, 546–55 (2019).

15. Ablasser, A., Goldeck, M., Cavlar, T., Deimling, T., Witte, G., Rohl, I., Hopfner, K.P., Ludwig, J., Hornung, V. cGAS produces a 2’-5’-linked cyclic dinucleotide second messenger that activates STING. Nature. 498, 380–4 (2013).

16. Ablasser, A., Schmid-Burgk, J.L., Hemmerling, I., Horvath, G.L., Schmidt, T., Latz, E., Hornung, V. Cell intrinsic immunity spreads to bystander cells via the intercellular transfer of cGAMP. Nature. 503, 530–4 (2013).

17. Moller, H.D., Mohiyuddin, M., Prada-Luengo, I., Sailani, M.R., Halling, J.F., Plomgaard, P., Maretty, L., Hansen, A.J., Snyder, M.P., Pilegaard, H., Lam, H.Y.K., Regenberg, B. Circular DNA elements of chromosomal origin are common in healthy human somatic tissue. Nat Commun. 9, 1069 (2018).

18. Wang, Y., Wang, M., Djekidel, M.N., Chen, H., Liu, D., Alt, F.W., Zhang, Y. eccDNAs are apoptotic products with high innate immunostimulatory activity. Nature. 599, 308–14 (2021).

19. Dillon, L.W., Kumar, P., Shibata, Y., Wang, Y.H., Willcox, S., Griffith, J.D., Pommier, Y., Takeda, S., Dutta, A. Production of Extrachromosomal MicroDNAs Is Linked to Mismatch Repair Pathways and Transcriptional Activity. Cell Rep. 11, 1749–59 (2015).

20. Wang, Y., Wang, M., Zhang, Y. Purification, full-length sequencing and genomic origin mapping of eccDNA. Nat Protoc. 18, 683–99 (2023).

21. Sankar, G., Karin, M. Missing Pieces in the NF-κB Puzzle. Cell. 109, S81–S96 (2002).

22. Henssen, A., MacArthur, I., Koche, R., H, D.-G. Purification and Sequencing of Large Circular DNA from Human Cells. Protocol Exchange. (2019).

23. Koche, R.P., Rodriguez-Fos, E., Helmsauer, K., Burkert, M., MacArthur, I.C., Maag, J., Chamorro, R., Munoz-Perez, N., Puiggros, M., Dorado Garcia, H., Bei, Y., Roefzaad, C., Bardinet, V., Szymansky, A., Winkler, A., Thole, T., Timme, N., Kasack, K., Fuchs, S., Klironomos, F., Thiessen, N., Blanc, E., Schmelz, K., Kunkele, A., Hundsdorfer, P., Rosswog, C., Theissen, J., Beule, D., Deubzer, H., Sauer, S., Toedling, J., Fischer, M., Hertwig, F., Schwarz, R.F., Eggert, A., Torrents, D., Schulte, J.H., Henssen, A.G. Extrachromosomal circular DNA drives oncogenic genome remodeling in neuroblastoma. Nat Genet. 52, 29–34 (2020).

24. Zhang, C.Z., Spektor, A., Cornils, H., Francis, J.M., Jackson, E.K., Liu, S., Meyerson, M., Pellman, D. Chromothripsis from DNA damage in micronuclei. Nature. 522, 179–84 (2015).

25. Stephens, P.J., Greenman, C.D., Fu, B., Yang, F., Bignell, G.R., Mudie, L.J., Pleasance, E.D., Lau, K.W., Beare, D., Stebbings, L.A., McLaren, S., Lin, M.L., McBride, D.J., Varela, I., Nik-Zainal, S., Leroy, C., Jia, M., Menzies, A., Butler, A.P., Teague, J.W., Quail, M.A., Burton, J., Swerdlow, H., Carter, N.P., Morsberger, L.A., Iacobuzio-Donahue, C., Follows, G.A., Green, A.R., Flanagan, A.M., Stratton, M.R., Futreal, P.A., Campbell, P.J. Massive genomic rearrangement acquired in a single catastrophic event during cancer development. Cell. 144, 27–40 (2011).

26. Toufektchan, E., Maciejowski, J. Purification of micronuclei from cultured cells by flow cytometry. STAR Protoc. 2, 100378 (2021).

27. Muller, K., Honcharova-Biletska, H., Koppe, C., Egger, M., Chan, L.K., Schneider, A.T., Kusgens, L., Bohm, F., Boege, Y., Healy, M.E., Schmitt, J., Comtesse, S., Castoldi, M., Preisinger, C., Szydlowska, M., Focaccia, E., Gaisa, N.T., Loosen, S.H., Jors, S., Tacke, F., Roderburg, C., Keitel, V., Bode, J.G., Boor, P., Davis, R.J., Longerich, T., Geisler, F., Heikenwalder, M., Weber, A., Vucur, M., Luedde, T. JNK signaling prevents biliary cyst formation through a CASPASE-8-dependent function of RIPK1 during aging. Proc Natl Acad Sci U S A. 118, (2021).

28. He, J., Gerstenlauer, M., Chan, L.K., Leithauser, F., Yeh, M.M., Wirth, T., Maier, H.J. Block of NF-kB signaling accelerates MYC-driven hepatocellular carcinogenesis and modifies the tumor phenotype towards combined hepatocellular cholangiocarcinoma. Cancer Lett. 458, 113–22 (2019).

29. Wolf, M.J., Adili, A., Piotrowitz, K., Abdullah, Z., Boege, Y., Stemmer, K., Ringelhan, M., Simonavicius, N., Egger, M., Wohlleber, D., Lorentzen, A., Einer, C., Schulz, S., Clavel, T., Protzer, U., Thiele, C., Zischka, H., Moch, H., Tschop, M., Tumanov, A.V., Haller, D., Unger, K., Karin, M., Kopf, M., Knolle, P., Weber, A., Heikenwalder, M. Metabolic activation of intrahepatic CD8+ T cells and NKT cells causes nonalcoholic steatohepatitis and liver cancer via cross-talk with hepatocytes. Cancer Cell. 26, 549–64 (2014).

30. de Almeida, T.M., Leitao, R.C., Andrade, J.D., Becak, W., Carrilho, F.J., Sonohara, S. Detection of micronuclei formation and nuclear anomalies in regenerative nodules of human cirrhotic livers and relationship to hepatocellular carcinoma. Cancer Genet Cytogenet. 150, 16–21 (2004).

31. Dungrawala, H., Rose, K.L., Bhat, K.P., Mohni, K.N., Glick, G.G., Couch, F.B., Cortez, D. The Replication Checkpoint Prevents Two Types of Fork Collapse without Regulating Replisome Stability. Mol Cell. 59, 998–1010 (2015).

32. Zhao, Y., Yu, L., Zhang, S., Su, X., Zhou, X. Extrachromosomal circular DNA: Current status and future prospects. Elife. 11, (2022).

33. Mohr, L., Toufektchan, E., von Morgen, P., Chu, K., Kapoor, A., Maciejowski, J. ER-directed TREX1 limits cGAS activation at micronuclei. Mol Cell. 81, 724–38 e9 (2021).

34. Sin, S.T., Deng, J., Ji, L., Yukawa, M., Chan, R.W., Volpi, S., Vaglio, A., Fenaroli, P., Bocca, P., Cheng, S.H., Wong, D.K., Lui, K.O., Jiang, P., Chan, K.C.A., Chiu, R.W., Lo, Y.M.D. Effects of nucleases on cell-free extrachromosomal circular DNA. JCI Insight. 7, (2022).

35. Pikarsky, E., Porat, R., Stein, I., Abramovitch, R., Amit, S., Kasem, S., Gutkovich-Pyest, E., Urieli-Shoval, S., Galun, E., Ben-Neriah, Y. NF-kappaB functions as a tumour promoter in inflammation-associated cancer. Nature. 431, 461–6 (2004).

36. Dou, Z., Ghosh, K., Vizioli, M.G., Zhu, J., Sen, P., Wangensteen, K.J., Simithy, J., Lan, Y., Lin, Y., Zhou, Z., Capell, B.C., Xu, C., Xu, M., Kieckhaefer, J.E., Jiang, T., Shoshkes-Carmel, M., Tanim, K., Barber, G.N., Seykora, J.T., Millar, S.E., Kaestner, K.H., Garcia, B.A., Adams, P.D., Berger, S.L. Cytoplasmic chromatin triggers inflammation in senescence and cancer. Nature. 550, 402–6 (2017).

37. Gluck, S., Guey, B., Gulen, M.F., Wolter, K., Kang, T.W., Schmacke, N.A., Bridgeman, A., Rehwinkel, J., Zender, L., Ablasser, A. Innate immune sensing of cytosolic chromatin fragments through cGAS promotes senescence. Nat Cell Biol. 19, 1061–70 (2017).

38. Yang, H., Wang, H., Ren, J., Chen, Q., Chen, Z.J. cGAS is essential for cellular senescence. Proc Natl Acad Sci U S A. 114, E4612–E20 (2017).

39. Liu, H., Zhang, H., Wu, X., Ma, D., Wu, J., Wang, L., Jiang, Y., Fei, Y., Zhu, C., Tan, R., Jungblut, P., Pei, G., Dorhoi, A., Yan, Q., Zhang, F., Zheng, R., Liu, S., Liang, H., Liu, Z., Yang, H., Chen, J., Wang, P., Tang, T., Peng, W., Hu, Z., Xu, Z., Huang, X., Wang, J., Li, H., Zhou, Y., Liu, F., Yan, D., Kaufmann, S.H.E., Chen, C., Mao, Z., Ge, B. Nuclear cGAS suppresses DNA repair and promotes tumorigenesis. Nature. 563, 131–6 (2018).

40. Zierhut, C., Yamaguchi, N., Paredes, M., Luo, J.D., Carroll, T., Funabiki, H. The Cytoplasmic DNA Sensor cGAS Promotes Mitotic Cell Death. Cell. 178, 302–15 e23 (2019).

41. Dunphy, G., Flannery, S.M., Almine, J.F., Connolly, D.J., Paulus, C., Jonsson, K.L., Jakobsen, M.R., Nevels, M.M., Bowie, A.G., Unterholzner, L. Non-canonical Activation of the DNA Sensing Adaptor STING by ATM and IFI16 Mediates NF-kappaB Signaling after Nuclear DNA Damage. Mol Cell. 71, 745–60 e5 (2018).

42. Charni-Natan, M., Goldstein, I. Protocol for Primary Mouse Hepatocyte Isolation. STAR Protoc. 1, 100086 (2020).

