## Supplementary figures, tables and detailed methods for "Extrachromosomal circular DNA promotes inflammation and hepatocellular carcinoma development"

### Figure S1

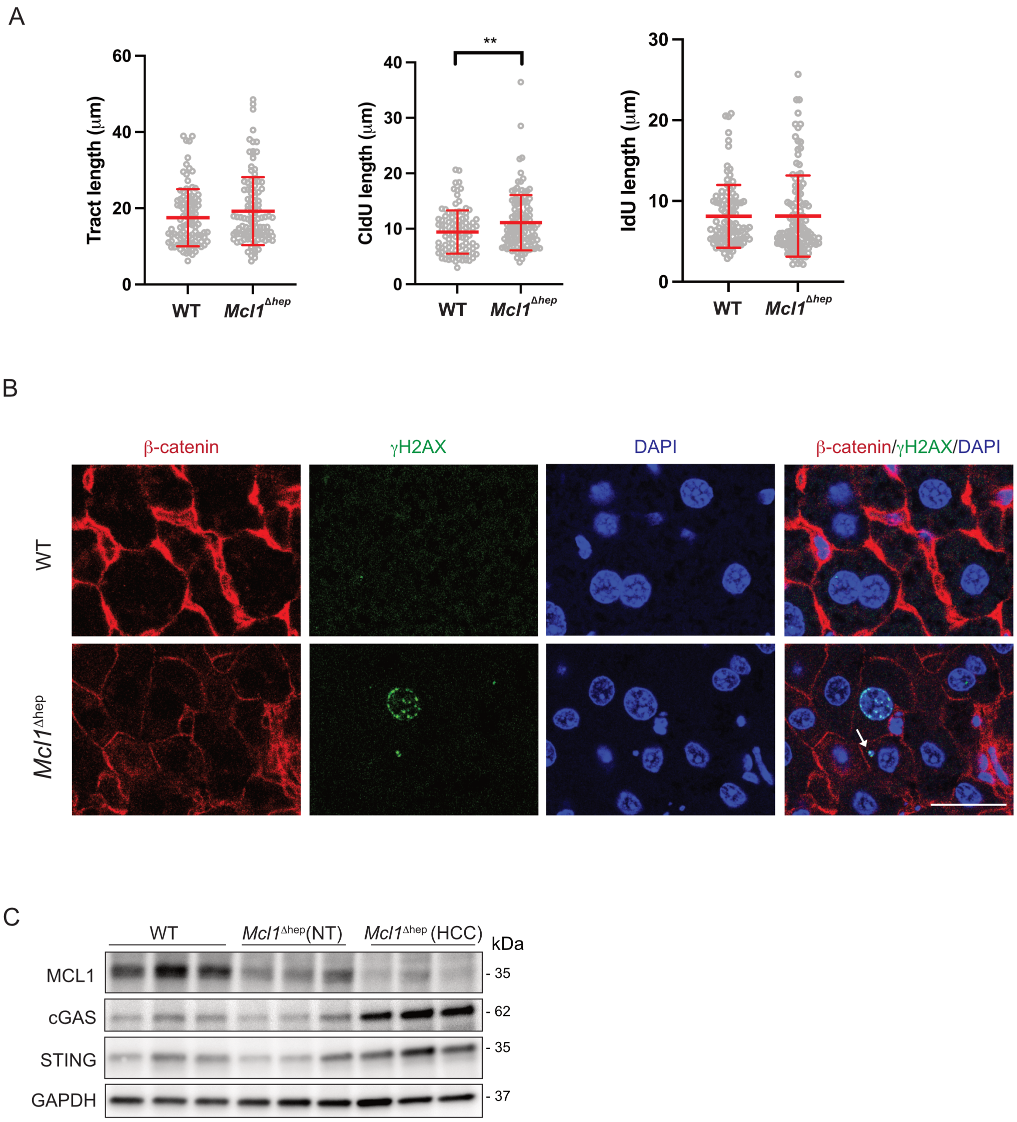


**Figure S1.** A) DNA fiber assay on primary hepatocytes from WT and *Mcl1*$^{\Delta hep}$ mice after 24 hours in culture. CldU: 5-chloro-2'deoxyuridine; IdU: 5-Iodo-2'-Deoxyuridine. Quantification of CldU track length, IdU track length, and CldU+IdU total track length. Student's t-test: ** $p<0.01$. B) Immunofluorescence staining indicating micronuclei (arrow) showing DNA damage in 12-month-old liver tissue. Scale bar: 25µm. C) Immunoblot of tissue extracts from the 12-month-old liver.

### Figure S2

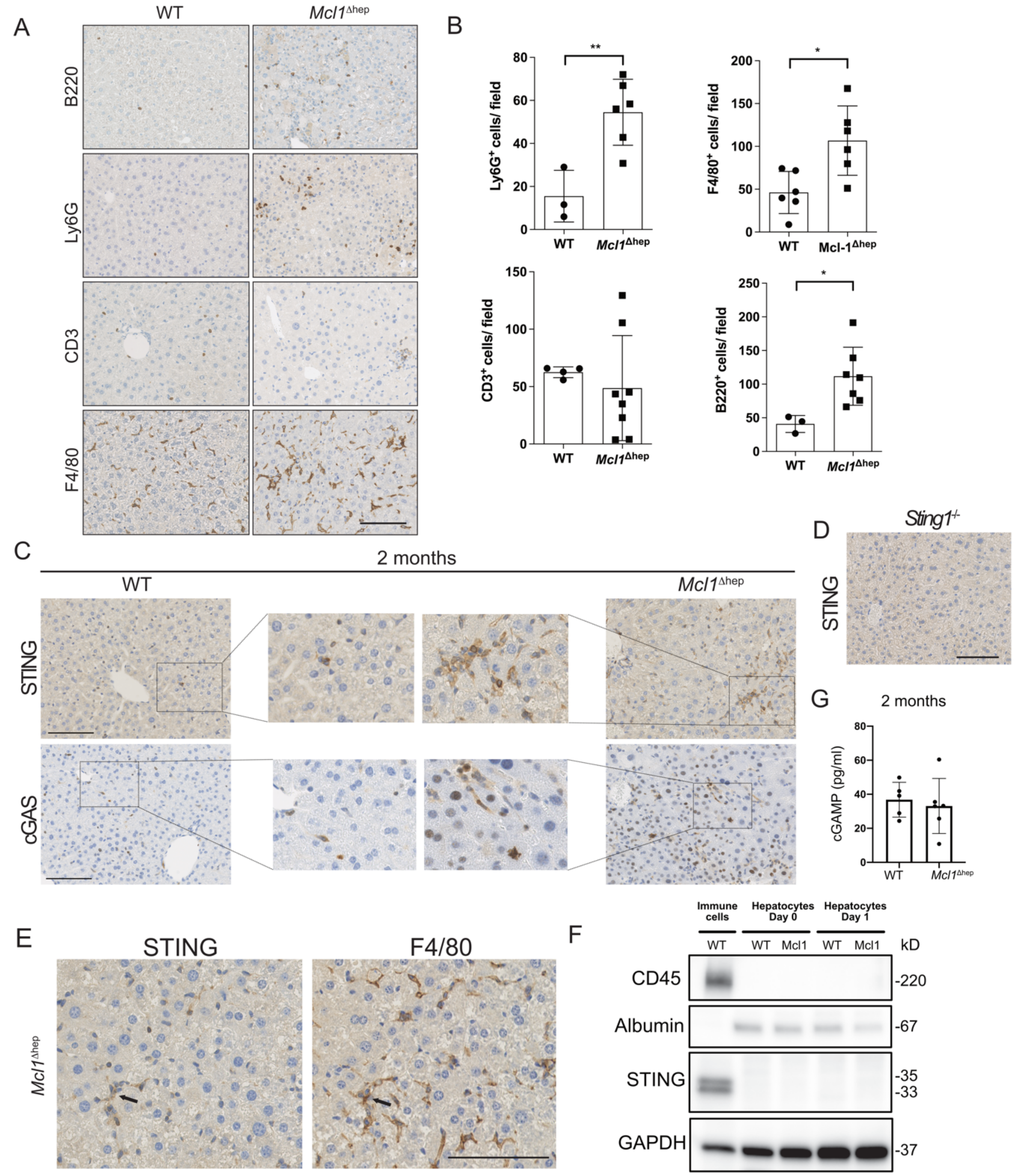


**Figure S2.** A) IHC staining of immune cell markers: B220, Ly6G, CD3, and F4/80. Scale bar: 50µm. B) Quantification of immune cells showing an increase in the infiltration of neutrophils, macrophages, and B cells. n≥3. Student's t-test: * $p<0.05$, ** $p<0.01$. C) IHC staining of STING and cGAS showing a different spatial expression pattern. Scale bar: 100µm. D) IHC staining of STING in *Sting1*$^{-/-}$ liver. Scale bar: 100µm. E) IHC staining of STING and F4/80 on serial sections showing the expression of STING and F4/80 in the same cells (arrow). Scale bar: 100µm. F)

Western blot from isolated immune cells and hepatocytes from mouse livers. WT: wild-type; Mcl1: *Mcl1*$^{\Delta hep}$. Immune cells and hepatocytes (Day 0) were used directly after isolation. Hepatocytes were then kept in culture for 24 hours (Day 1) and protein was isolated. G) Measurement of cGAMP level from protein lysates using ELISA. n≥5.

### Figure S3

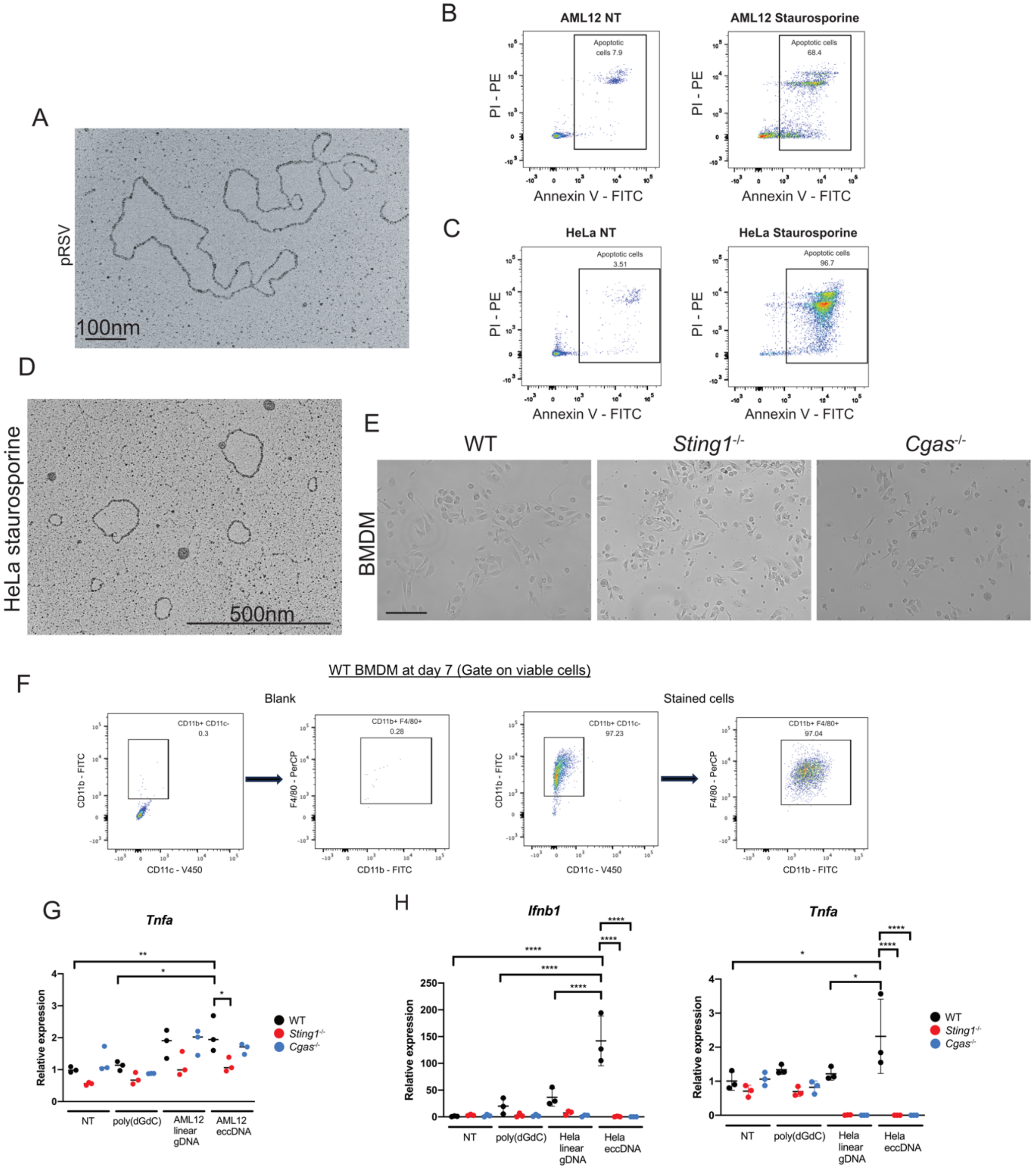


**Figure S3.** A) Electron micrograph of plasmid pRSV. Scale bar: 100nm. B&C) Flow cytometry analysis of AML12 and HeLa cells treated with staurosporine for 24 hours. Cells were stained with Annexin V and PI for labelling early and late apoptotic cells. D) Visualization of eccDNA isolated from HeLa cells treated with 0.5M of staurosporine for 24 hours. Scale bar: 500nm. E) BMDM differentiated from bone marrow cells stimulated with M-CSF for 7 days. Scale bar: 50μm F) Phenotyping of WT BMDM by flow cytometry. G) BMDMs from WT, *Sting1*$^{-/-}$ and *Cgas*$^{-/-}$ mice were transfected with 10ng/ml of eccDNA, linear genomic DNA, and poly(dGdC). The *Tnfa* expression was analyzed 12 hours after the transfection. n=3. H) BMDM from WT, STING-

/- and cGAS-/- mice were transfected with 100ng/ml of poly(dGdC), linear gDNA from HeLa cells, or eccDNA from HeLa cells. The expression of *Ifnb1* and *Tnfa* were detected by qPCR. n=3. One-way ANOVA: * $p<0.05$, ** $p<0.01$, *** $p<0.001$, **** $p<0.0001$.

**Figure S4**

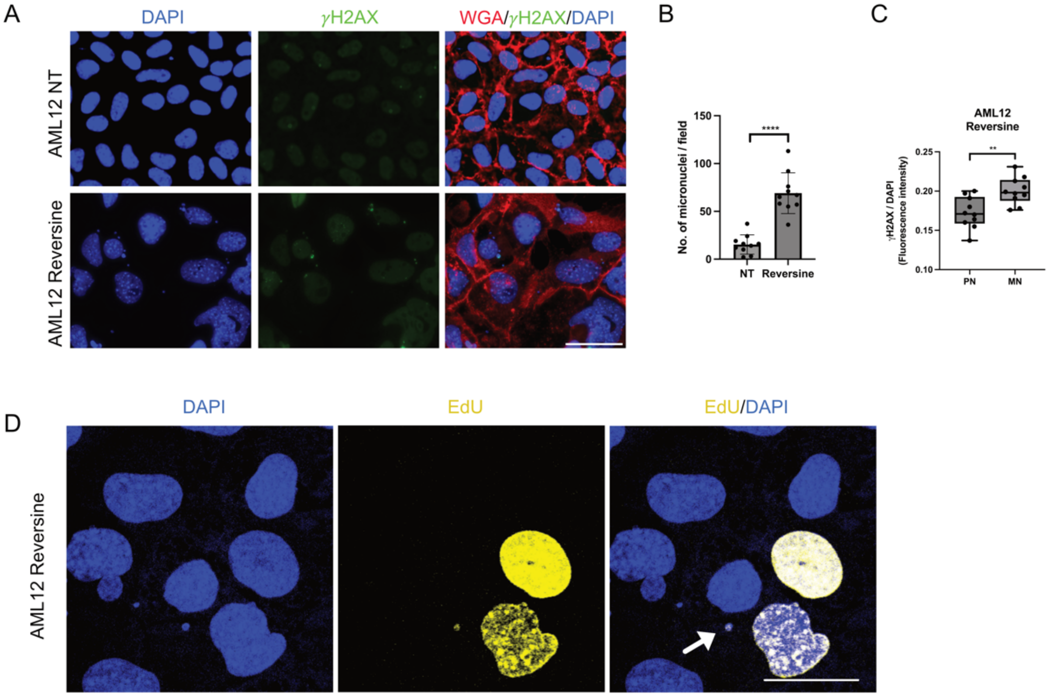


**Figure S4.** A) AML12 cells treated with reversine for 48 hours showed an increase in micronuclei levels. NT: non-treated. WGA: wheat germ agglutinin (membrane dye). Scale bar: 40μm. B) Quantification of the number of micronuclei observed per captured field. Student's t-test: **** $p<0.0001$. C) Comparison of γH2AX intensity normalized to DAPI intensity between primary nuclei (PN) and micronuclei (MN). Student's t-test: ** $p<0.01$. D) AML12 cells were labeled with EdU for 2 hours at the end of the reversine treatment. Arrow indicated a S phase micronuclei. Scale bar: 20μm.

### Figure S5

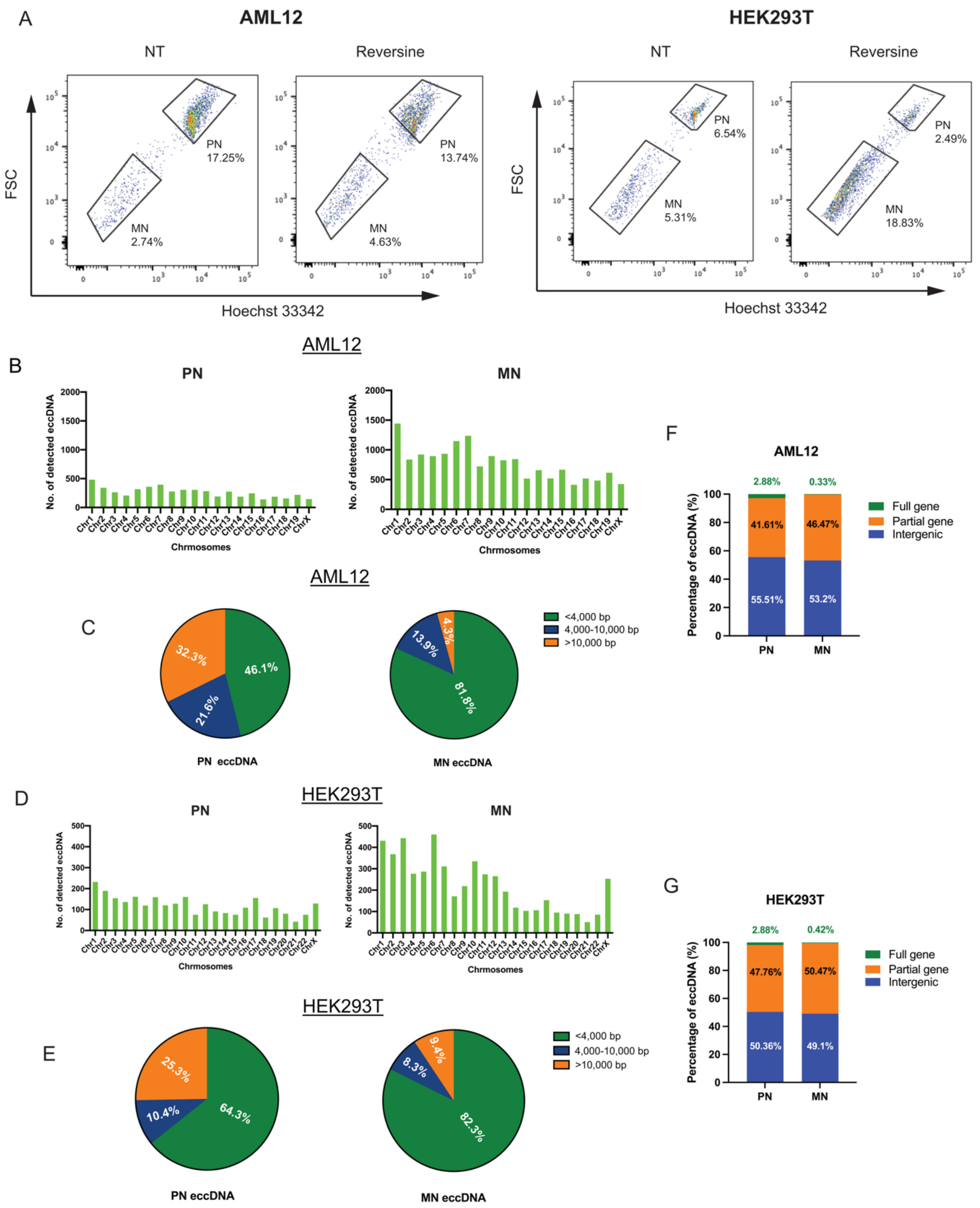


**Figure S5.** A) FACS sorting of primary nuclei and micronuclei from AML12 cells and HEK293T cells 48 hours after reversine treatment. B&D) eccDNA detected by cirSeq from isolated primary nuclei (PN) and micronuclei (MN) were mapped to the genome of mouse (AML12) and human (HEK293T) and the frequency of detected eccDNA was plotted across all chromosomes. C&E) Pie charts showing the percentage of eccDNA with size <4,000bp, 4,000-10,000bp, and

>10,000bp. F&G) Bar charts showing the percentages of eccDNA containing full gene, partial gene and intergenic sequences in PN and MN.

**Figure S6**

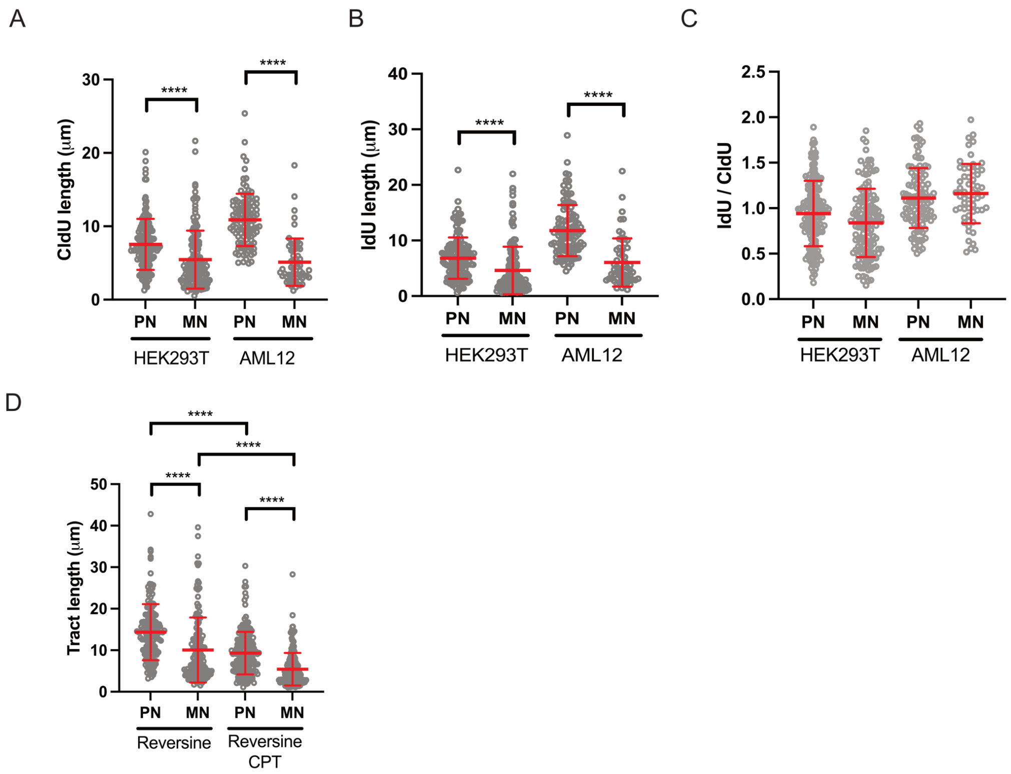


**Figure S6.** A-C) NuSeF assay comparing the DNA fibers from primary nuclei (PN) and micronuclei (MN) of HEK293T and AML12 cells. CldU and IdU were pulse-labelled for 30 minutes each. The CldU tract length, IdU tract length, or IdU/CldU ratio was compared between PN and MN. Student's t-test: **** $p<0.0001$. D) Tract length (CldU + IdU) measurement from DNA fibers obtained from PN and MN from HEK293T cell treated with reversine and with/without CPT. One-way ANOVA: **** $p<0.0001$.

### Figure S7

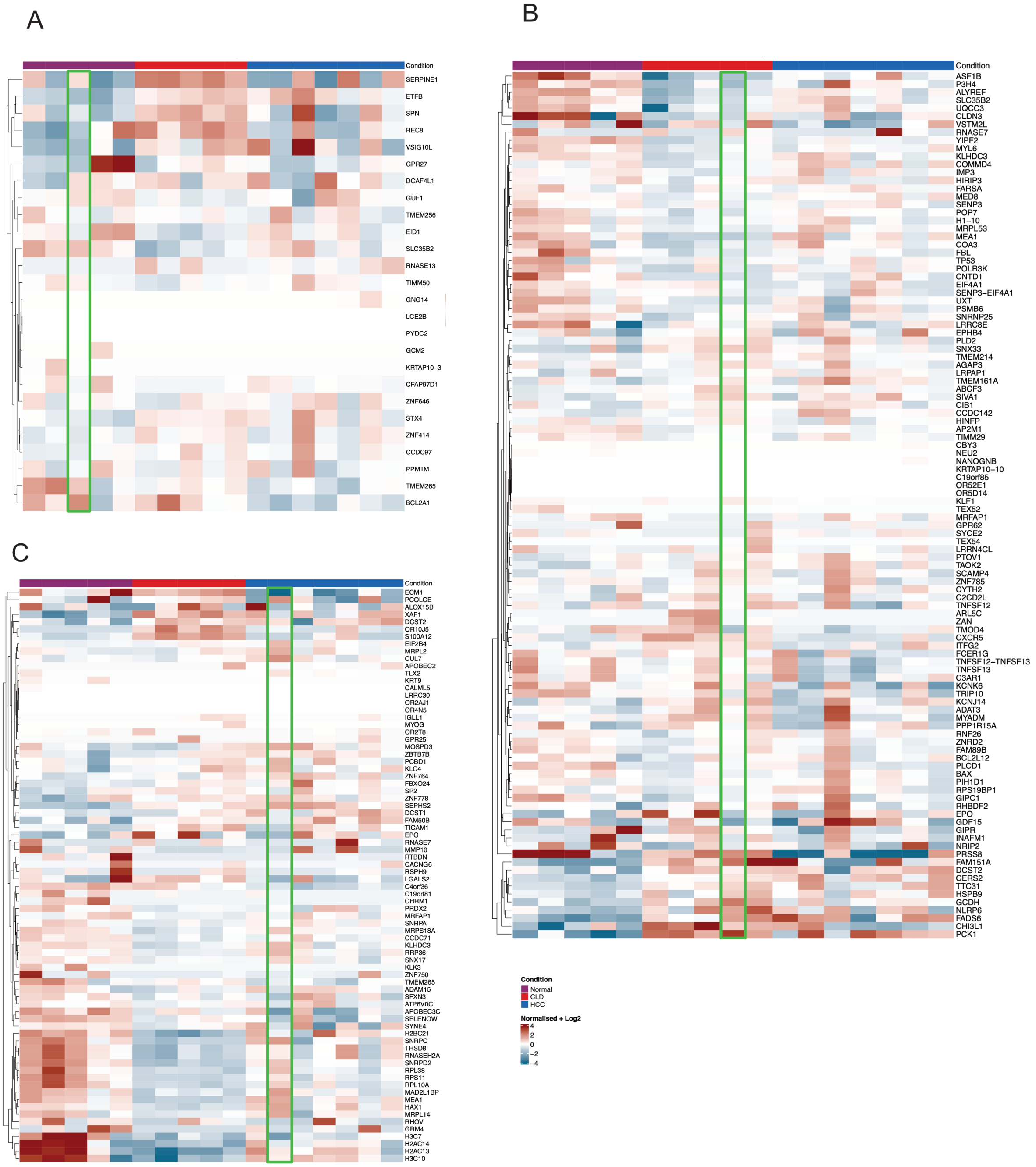


**Figure S7.** Heatmap showing the expression of genes across normal, CLD, and HCC samples. Gene lists shown were genes found in eccDNA of A) normal, B) CLD, and C) HCC samples. The samples with the list of genes present in eccDNA were highlighted with green rectangles.

### Figure S8

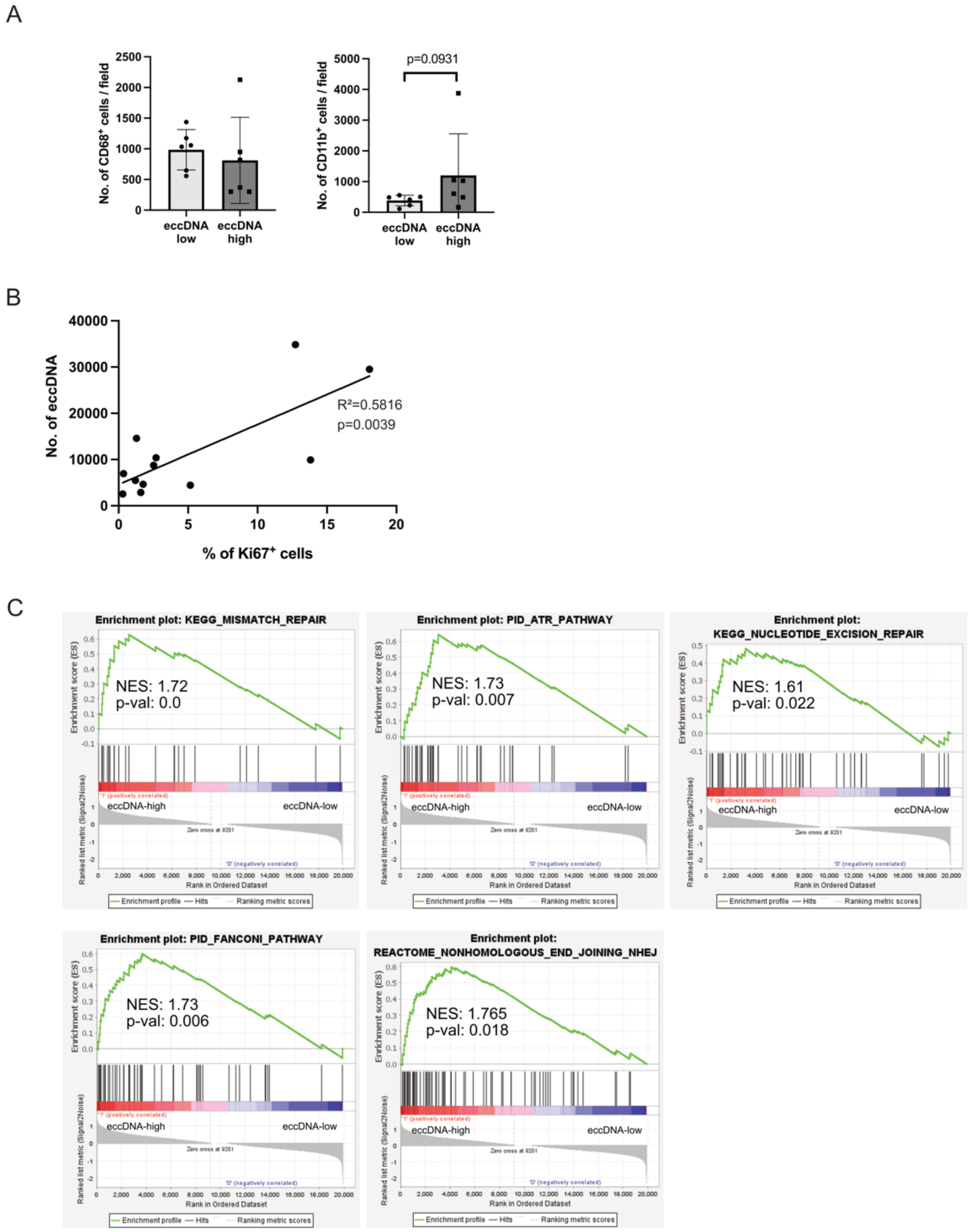


**Figure S8.** A) Comparison of $CD68^+$ and $CD11b^+$ cells between eccDNA-high and eccDNA-low groups. Mann-Whitney test. B) Correlation of percentage of $Ki67^+$ cells with the number of detected eccDNA in CLD and HCC samples. C) GSEA analysis showing enrichment gene signatures related to DDR and DNA repair pathways.

**Figure S9**

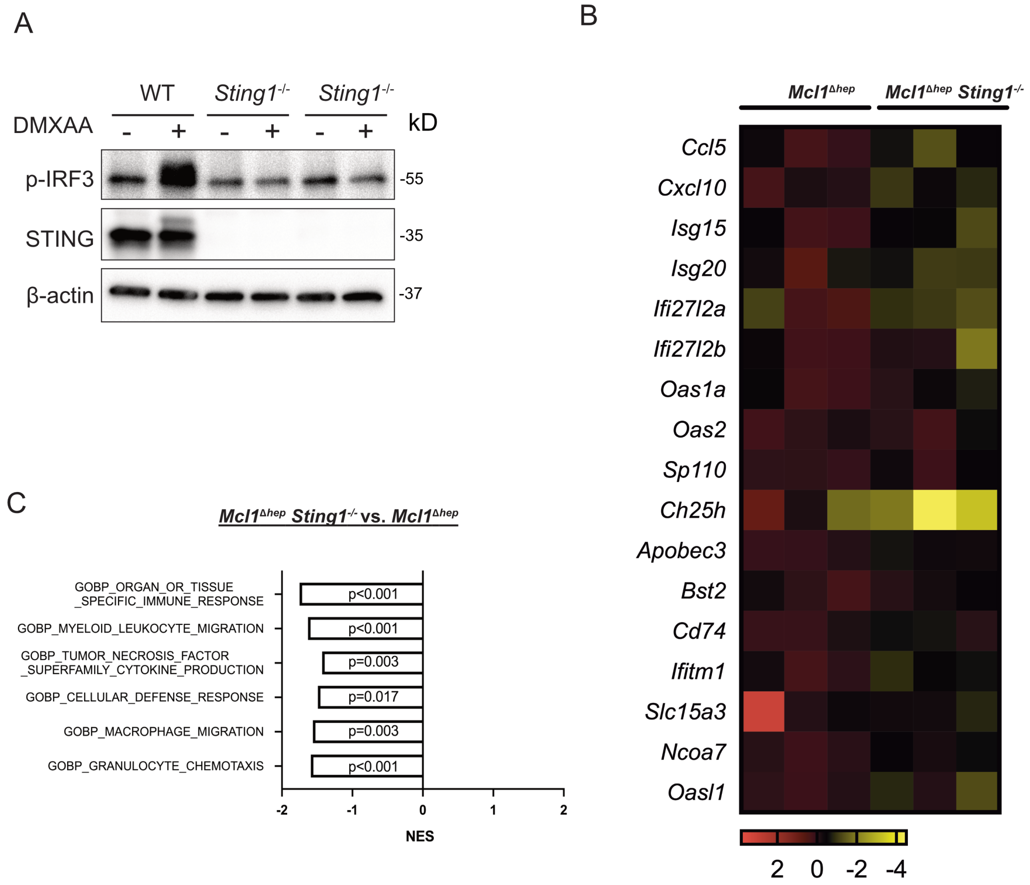


**Figure S9.** A) Western blot from splenocytes obtained from WT and *Sting1*$^{-/-}$ mice. Splenocytes were treated with or without DMXAA for 3 hours. B) Heatmap of ISGs from RNAseq data. C) Depleted GSEA signatures in *Mcl1*$^{\Delta hep}$ *Sting1*$^{-/-}$ mice.

**Extended Methods**

Cell lines

AML12 (CRL-2254), HEK293T (CRL-3216), and HeLa (CCL-2) cells were obtained from ATCC. Mycoplasma, bacterial, and fungal testing was performed by the supplier with the information that none of these contaminants were detected.

Serum enzyme activities measurement

To determine the serum ALT and AST activities, approximately 50 µl of blood was collected from the tail vein of mice at the indicated time points using BD Microtainer SST Tubes (BD #365968). After settling at room temperature for at least 10 minutes, serum was obtained by centrifugation at 8500g for 90 seconds. ALT and AST activities were measured using Reflotron Plus (Roche) with GOT/AST and GPT/ALT stripes (Roche #10745120202 & 10745138202).

Histology, immunohistochemistry (IHC), immunofluorescence (IF) and microscopy

For histological analysis, mice were euthanized and livers were harvested and fixed in 4% neutral buffered formalin (PFA) for at least 24 hours. Liver tissues were then dehydrated and embedded with paraffin. Paraffin sections of 3µm thickness were used in IHC and IF staining.

Hematoxylin and eosin (H&E) were performed by the standard protocol. For IHC, immune stains were performed by Leica Bond RX. Stained IHC slides were scanned by Hamamatsu C9600 slide scanner. For IF, stains were performed manually, using heat-mediated antigen retrieval (pressure cooker, 10 minutes) with citrate buffer (pH 6) after deparaffinization and rehydration. The tissue sections were blocked with 5% BSA/PBS solution for one hour at room temperature before overnight primary antibody incubation followed by one hour of fluorophore-tagged secondary antibody incubation at room temperature. The slides were mounted with EverBrite hardset mounting medium carrying DAPI (Biotium #23004) for microscopy. IF images were acquired with Leica DM6 microscope or Leica SP8 confocal microscope. Images were analyzed using QuPath (version 0.4.4) or ImageJ (Fiji). A list of IHC and IF antibodies is provided in Table S2.

Western blot

Liver tissues were harvested and snap-frozen followed by homogenization through powderization. Total proteins were extracted with 4% sodium dodecyl sulfate (SDS)/100mM Tris-HCl. Protein concentration was determined by the BCA Protein Assay Kit (Thermo Scientific #23228). For Western blots, protein lysates (10-20μg for tissue lysates; 5μg for cell lysates) were prepared with NuPage LDS Sample Buffer (Invitrogen #NP0007) and 4-20% mini-PROTREAN TGX gels (Bio-rad #4568096 and #4568094) were used to separate proteins. Protein was transferred by the Transblot-turbo system (Bio-Rad) onto PVDF membranes. The membranes were blocked with 5% BSA in TBS buffer/0.1% Tween-20 or 1X ROTI Block (ROTH #A151.2) for 1 hour. Primary and secondary antibodies were then applied. The band signal was detected by Fushion Solo. Densitometry of the Western blot bands was performed using ImageJ. A list of Western blot antibodies is provided in Table S3.

Primary hepatocyte and immune cell isolation from the liver

Hepatocyte isolation was performed according to the proposed protocol (1). Briefly, mice were first anesthetized by ketamine (100mg/kg) and xylazine (16mg/kg). A pre-warmed (37°C) pre-perfusion solution (0.5mM EDTA/20mM HEPES/HBSS) was perfused via the vena cava using a constant flow rate of 7ml/min regulated by a peristaltic pump (VWR). Once yellow spots were seen in the liver, the portal vein was cut to release the blood and perfusion buffer. The liver was first perfused with the pre-perfusion buffer for 5 minutes and then switched to perfusion solution (20mM HEPES / 1X Penicillin/Streptomycin / 3mM $CaCl_2$ / DMEM/F12 / 0.2mg/ml Liberase (Roche #05401127001)) for 10 minutes. At the end of the liberase digestion, the liver was carefully transferred to a 10cm petri dish and washed with Wash solution (4% FBS / 1X Penicillin/Streptomycin / Willian's E medium) and the gall bladder was removed. The liver was then transferred to a new petri dish with 10ml of Wash solution and hepatocytes were gently released from the liver using forceps. The Wash solution containing cells was filtered through a 70μm cell strainer (Falcon #352350) and stored on ice. The cell suspension was centrifuged at 20g for 3 minutes. The pellet was used for hepatocyte isolation and the supernatant, which contained non-parenchymal cells, was transferred to a new 50ml Falcon tube for immune cell isolation. For hepatocyte purification, the pellet was resuspended with 10ml of Wash solution and mixed with 10ml of 90% percoll solution (Cytiva #GE-17-0891-01) and centrifuged at 200g for 10 minutes. This step was repeated one more time to increase the purity of viable cells. The pellet was then resuspended in Wash solution and seeded at a density of $3x10^5$ cells/well in a collagen pre-coated 24-well plate. The medium was exchanged to Hepatocyte medium (Wash solution / 50ng/ml EGF / 1μg/ml insulin / 10μg/ml Transferrin / 1.3μg/ml Hydrocortisone) after 4 hours. A lower seeding density was used for hepatocytes in the DNA fiber assay. For isolating immune cells, the volume was brought to 50ml using the Wash solution. The solution was centrifuged at 20g for 3 minutes and the supernatant was transferred to a new 50ml Falcon tube. This step was repeated for additional two times. The transferred supernatant was then centrifuged at 2000 rpm for 5 minutes. The immune cell pellet was resuspended in 10ml of 36% Percoll solution and centrifuged at

2000rpm for 20 minutes at 4°C. RBCs were lysed with 1X RBC lysis buffer (G Biosciences #786-649) at room temperature for 5 minutes. The immune cells were washed once with 10ml of PBS and pelleted followed by snap-freezing and storage.

Preparation of bone-marrow-derived macrophages (BMDM)

BMDM was prepared according to the proposed protocol (2). Briefly, mice were euthanized using $CO_2$. The femur and tibia from both limbs were collected in 1.5ml Eppendorf tubes and stored on ice. Under a cell culture hood, 1ml of cold PBS was carefully injected through the collected femur and tibia to flush out the bone marrow cells into a well of a 6-well plate. The collected bone marrow cells were centrifuged at 200g for 5 minutes at 4°C. The cell pellet was resuspended in 1X RBC lysis buffer and incubated at room temperature for 5 minutes to remove erythrocytes. The cells were then washed once with 10ml of cold PBS and pelleted. The cell pellet was then resuspended in 12 ml of Bone Marrow Culture Medium (DMEM / 10% FBS/ 1% Penicillin/Streptomycin / 10ng/ml M-CSF) and distributed into a 6-well plate, with 2ml per well. Each well carried approximately 3 million cells. On day 4, half of the medium was exchanged with fresh Bone Marrow Culture Medium. BMDMs were ready for experiments on day 7.

Transfection of poly(dGdC), linear gDNA and eccDNA

For generating linear gDNA, total gDNA was isolated from AML12 cells and HeLa cells. An aliquot of the gDNA (1µg) was sonicated using Bandelin SONOPLUS Mini 20 for 50 seconds. The size of DNA fragments was confirmed to be below 3kb by gel electrophoresis. Transfections of poly(dGdC) (InvivoGen #tlrl-pgcn), linear gDNA, and eccDNA were performed by Lipofectamine 3000 Transfection Reagent (Invitrogen #L3000008) according to the manufacturer's instruction. Transfected BMDMs were collected 12 hours after the transfection. Controls received only transfection reagents.

Flow cytometry

For detection of apoptotic cells, cells were treated with staurosporine (1μM) (MedChemExpress #HY-15141) for 24 hours or reversine (0.5μM) (MedChemExpress # HY-14711) for 48 hours. Cells were trypsinized and washed twice with cold PBS. Pelleted cells were resuspended in Annexin V binding buffer (BioLegend #640914). Approximately 1 million cells in a volume of 100μl were prepared in a 5ml test tube. Annexin V-FITC and Propidium Iodine were added to the cells followed by an incubation for 15 minutes at room temperature in the dark. At the end of incubation, 400μl of Annexin V binding buffer was added to the tube, and the cells were analyzed with BD FACS Canto II. A list of Western blot antibodies is provided in Table S4.

Quantitative real-time PCR (qPCR)

Total RNA was isolated from pulverized liver tissues or cells using the RNeasy mini kit or RNeasy micro kit (Qiagen #74004 and #74104). Complementary DNA (cDNA) synthesis was performed using a High Capacity cDNA Reverse Transcription Kit (Applied Biosystems #4368814) with the extracted mRNA from each sample. qPCR was performed using SYBR Green PCR master mix (Applied Biosystem #43-687-02) with ViiA7 Real-Time PCR system (Applied Biosystems). A list of primers is provided in Table S5.

Primary nuclei and micronuclei isolation

Purification of micronuclei and primary nuclei was performed following a published protocol (3). Briefly, AML12 (6 x 150mm plates) and HEK293T cells (3 x 150mm plates) were treated with reversine for 48 hours to promote micronuclei formation. On the day of the experiment, cells were harvested and incubated with cytochalasin B for 30 minutes at 37°C. The cells were pelleted and resuspended in 5 ml of lysis buffer. The cell lysate was then mixed with an equal volume of 1.8M sucrose buffer. A sucrose gradient was prepared by first adding 15 ml of 1.6M sucrose buffer at the bottom of a 50 ml falcon tube followed by layering 20 ml of 1.8M sucrose buffer on top. The cell lysate/1.8M sucrose buffer mixture was laid on top of the prepared sucrose gradient and centrifuged at 950g for 20 minutes at 4°C. The top 2ml was discarded. The next 5 ml of the micronuclei-enriched fraction was transferred to another 50 ml falcon tube and washed with 20 ml of PBS supplemented with 1X protease inhibitor followed by centrifugation at 1500g for 20 minutes at 4°C. The upper supernatant was discarded, leaving only 1 ml for resuspending the pellet. The sample was filtered through the cell strainer of the FACS round-bottom tube (Falcon #352235). Hoechst 33342 (1μg/ml) (Life technology #H3570) and MitoView Green (Biotium #70054-T) (40nM) were added to the sample 10 minutes before FACS sorting. Primary nuclei and micronuclei were sorted according to the size and Hoechst 33342 intensity using BD FACSAria III. MitoView dye was used to minimize mitochondria contamination during sorting.

<u>eccDNA isolation and CirSeq</u>

We used the eccDNA isolation protocol proposed by Wang et al., 2023 for purifying eccDNA for transfection experiments and electron microscopy (4). Briefly, AML12 cells or HeLa cells were treated with 1μM staurosporine for 24 hours or with 0.5μM reversine for 48 hours. Cells were collected and washed twice with PBS. The cells were pelleted and resuspended in 10ml of suspension buffer (10mM EDTA pH8 / 150 mM NaCl / 1% glycerol / lysis blue / RNase A / β-mercaptoethanol). 10 ml of Pyr buffer (0.5M pyrrolidine / 20mM EDTA / 1% SDS / β-mercaptoethanol / pH 11.8) was added to the resuspended cells and gently mixed well. The mixture was incubated at room temperature for 5 minutes. 10 ml of Buffer S3 (Qiagen Plasmid Plus Midi kit #12943) was then added to the solution and mixed well until the solution turned white. The mixture was centrifuged at 3148g for 20 minutes at 4°C. The clear lysate was filtered through the QIAfilter Cartridge (Qiagen Plasmid Plus Midi kit) and mixed with 1/3 volume of Buffer BB. Using QIAvac 24 plus, all lysates were passed through the QIAGEN Plasmid Plus spin column. The spin columns were then washed with ETR buffer and PE buffer. Crude circular DNA extract was eluted with 100μl of DNase-free water. The crude circular DNA concentration was determined by Qubit 1X dsDNA HS Assay Kit (invitrogen #Q33230). Next, plasmid-safe DNase (Lucigen #E3101K) with a final concentration of 0.4U/μl (1X Plasmid-safe Reaction Buffer / 1mM ATP / 10U Plasmid-safe DNase) was mixed with 3μg of crude circular DNA to eliminate the contaminating linear DNA. PacI was added to facilitate the elimination of mitochondrial DNA. The digested DNA was then cleaned up using the phenol/chloroform/isoamyl alcohol method. DNA was then precipitated from the aqueous fraction by adding 1μl of glycogen, 1/10 volume of sodium acetate (3M, pH5.5), and 3 volumes of 200 proof ethanol with incubation at -80°C for at least 3 hours. The precipitated DNA was centrifuged at 20,000g for 30 minutes at 4°C. The DNA pellet was washed once with 1 ml of freshly prepared 80% ethanol and resuspended in 50 μl of 2 mM Tris-HCl (pH 7). eccDNA was then selectively purified using Solution A (Bingene #220501). DNA resuspended in 2mM of Tris-HCl was mixed with 700μl of Solution A and incubated at room temperature for 5 minutes. The solution was then mixed well with 10μl of Dynabeads MyOne Silane beads (Invitrogen #37002D) and put on a magnetic holder (Invitrogen DynaMag-2

#12321D). After settling for 2 minutes, all solution was discarded without disturbing the beads. The beads were washed twice with 300µl of Solution A using the same procedure. While leaving the tubes on the magnetic holder, the beads were washed twice with 700µl of 3.5M NaCl and twice with 800µl of freshly prepared 80% ethanol. eccDNA was eluted with 20µl of 0.1X elution buffer (Qiagen Plasmid Plus Midi kit).

For the purification of eccDNA from tissue, the isolation of crude circular DNA was performed using the QIAprep Spin Miniprep Kit (Qiagen # 27104). 50 mg of frozen tissue obtained from animals were cut into small pieces and digested overnight with proteinase K at 0.6U/µl (Thermo #EO0491) in 600µl of P1 solution (QIAprep Spin Miniprep Kit # 27104) at 50°C with shaking at 650rpm. The lysate was centrifuged at 1,000 rpm for 2 minutes and the supernatant was transferred to a 2ml eppendorf tube (DNase free) and LyseBlue was added at 1:1000 to the collected supernatant. 600µl of Buffer P2 was then added to the lysate and incubated at room temperature for 5 minutes. Immediately after the incubation, 840µl of Buffer N3 was added to neutralize the solution. The mixed solution was centrifuged at 13,000 rpm for 10 minutes at room temperature. 800µl of the supernatant was transferred to a QIAprep 2.0 spin column. The column was centrifuged for 30 seconds and the flow through was discarded. The column was then washed with 0.5ml of Buffer PB followed by 0.75ml of Buffer PE. The column was then dried by centrifuging at a maximum speed for 1 minute and the DNA was eluted with 100µl of DNase-free water. eccDNA purification was then performed as described above.

For preparing DNA samples for cirSeq, we used a separate protocol that was suggested to preserve large circular DNA and reduce eccDNA loss due to the potential linearization of eccDNA by restriction enzymes (5). Briefly, high-molecular-weight (HMW) DNA was isolated from tissue using the MagAttract HMW DNA kit (Qiagen #67563) according to the manual of the manufacturer. Around 5-10 mg of tissue was obtained and briefly washed with cold PBS. The tissue was briefly spanned down and resuspended in 220µl of Buffer ATL. 20µl of proteinase K was added to the sample and mixed well by vortexing. The sample was digested overnight at 56°C with shaking at 900rpm. After digestion, 200µl of the samples were transferred to a new 2ml Eppendorf tube. For each sample, 4µl of RNase A was added and incubated for 2 minutes at room temperature. Then, 150µl of Buffer AL 280µl of Buffer MB and 40µl of MagAttact Suspension G were added to each sample and incubated for 3 minutes at room temperature. The tubes were transferred to a Magnetic rack and waited for 1 minute until the beads were completely separated. The beads were washed twice with 700µl of Buffer MW1 by resuspending the beads and incubating at room temperature for 2 minutes followed by removal of the buffer using the magnetic rack. The beads were then washed twice with 700µl of Buffer PE. While keeping on the magnetic holder, the beads were then rinsed twice with 700µl of DNase-free water. The tubes were removed from the magnetic holder and the HMW DNA was eluted with 150µl of DNase-free water with shaking at 1400rpm for 3 minutes at room temperature. The purified HMW DNA was obtained by separating the beads from the elute using the magnetic holder. The eluted HMW DNA was transferred to a new 1.5ml DNA low-bind tube. The HMW DNA concentration was determined by Qubit dsDNA HS Assay Kit (Invitrogen #Q32850).

Digestion of linear DNA was performed using plasmid-safe DNase (Lucigen #E3101K) with a final concentration of 0.2U/µl (1X Plasmid-safe Reaction Buffer / 1mM ATP / 20U Plasmid-safe DNase) for every 5µg of HMW DNA for 5 days at 37°C. For PN and MN samples, 50ng of DNA was used in this step. The Plasmid-safe DNase and ATP were replenished every 24 hours. At the end of the 5-day digestion cycle, the DNase was inactivated at 70°C for 30 minutes. Rolling circle amplification was performed using the REPLI-g Mini Kit (Qiagen #150025) to amplify the circular DNA in the plasmid-safe DNase digested sample. In a 0.2ml PCR tube, 5µl of the digested HMW DNA was mixed with 5µl of Buffer D1 and incubated at 25°C for 3 minutes. After that, 10µl of Buffer N1 was added to the sample and mixed well by vortexing. 30µl of REPLI-g DNA polymerase master mix (29µl of REPLI-g Mini Reaction Buffer and 1µl of REPLI-g Mini DNA polymerase) were then added to each sample and incubated at 30°C for 16 hours. The sample was then inactivated at 65°C for 3 minutes. The quantity of double-stranded DNA was measured by Qubit BR dsDNA kit (Invitrogen #Q32850).

Library Preparation was performed on a 96-well plate using the NEBNext® Ultra™ II FS DNA Library Prep Kit for Illumina (NEB # E7805L). 20 ng of the RCA product from each sample was fragmented with 1.75µl of Ultra II FS Reaction Buffer and 0.5µl of Ultra II FS Enzyme Mix in a thermal cycler at 37°C for 25 minutes and 65°C for 30 minutes. Adaptor ligation was performed with 1µl of Adaptor (0.04 µl of adaptor diluted by 0.96 µl of 10 mM Tris-HCl, pH 7.5-8.0 with 10 mM NaCl), 7.5 µl of Ligation Master Mix and 0.25 µl of Ligation Enhancer. After 15 minutes of incubation at 20°C, 1 µl of diluted USER (0.75µl of USER with 0.25µl of Nuclease-free water) was added and the samples were incubated at 37°C for 15 minutes. Adaptor-ligated DNA was cleaned with 0.8X (14.8 µl) of AMPure XP Beads from Beckman Coulter (A63881), mixed and incubated for 5 minutes. The samples were kept on a magnetic holder (Invitrogen DynaMag™-96 Magnet # 12027). The supernatant was discarded and the beads were washed twice with 100 µl of 80 % Ethanol while the plate was on the magnet. All ethanol was discarded and the beads were dried for 2 minutes. DNA was eluted with 6µl of Elution Buffer (Qiagen Buffer EB #19086). 5µL of the eluted DNA from each well was transferred into a new 96-well plate.

PCR enrichment of 5µL adaptor-ligated DNA was performed by adding 2.5 µl of pre-mixed unique dual index primer pairs (NEBNext 96 unique dual index primer pairs kit 3 NEB #E6444L). Each well must contain a different primer pair for multiplexing. 6.25 µL of Ultra II Q5 Master Mix was added to each well. In a thermocycler, run a PCR program: 98°C for 30 seconds, 98°C for 10 seconds and 65°C for 75 seconds for 7 cycles and finish with 65°C for 5 minutes. The PCR reaction was cleaned using 10 µl (0.8X) of AMPure XP Beads from Beckman Coulter (A63881), mixed and incubated for 5 minutes. The supernatant was discarded without touching the beads and the beads were washed twice with 100 µl of 80 % Ethanol while the plate was on the magnetic holder (Invitrogen DynaMag™-96 Magnet # 12027). The beads were dried for 2 minutes and DNA was eluted in 30 µl of Elution Buffer (Qiagen Buffer EB #19086) after 5 minutes of incubation. 28 µl of supernatant from each well was transferred to a new 96-well plate. The concentration of all samples was determined using Qubit 1X dsDNA HS Assay-Kit (ThermoFisher scientific # Q33231). Quality control of libraries was performed using TapeStation (Agilent High Sensitivity D1000 Reagents # 5067-5585 and High Sensitivity D1000 ScreenTape # 5067-5584). All samples were brought to the same concentration and 3 µl of each row of the 96-well plate was pooled into a 0.2 ml 8-tube strip. 10µl of each 0.2 ml tube was pooled into a 1.5 ml DNA LoBind tube. A cleanup step was performed with 64 µl (0.8X) of AMPure XP Beads in 80 µl of the pooled libraries using a magnetic holder (DynaMag™-2 Magnet # 12321D). The beads were washed twice with 200 µl of 80 % ethanol using the magnetic holder. After drying for 2 minutes, samples were eluted in 20 µl of Elution Buffer (Qiagen Buffer EB #19086). 18µl of the eluted supernatant was transferred to a new DNA LoBind tube. The concentration of all samples was determined using Qubit 1X dsDNA HS Assay-Kit (ThermoFisher scientific # Q33231). A quality control step of libraries was performed again using TapeStation (Agilent High Sensitivity D1000 Reagents # 5067-5585 and High Sensitivity D1000 ScreenTape # 5067-5584). The DNA libraries were submitted for 150bp pair-end Illumina sequencing using NovaSeq X Plus 10B.

To detect the presence of eccDNA, adapter sequences were trimmed from the reads using Trim Galore v. 0.6.10. The reads were then aligned either to the human reference genome hg38 or the mouse reference genome mm39 using the Burrows–Wheeler Aligner MEM v. 0.7.17 with default parameters. PCR and optical duplicates were removed with biobambam2 v. 2.0.183. The resulting BAM files were then analyzed to identify split reads and outward-facing discordant read pairs, which indicate circle-supporting reads. The coordinates of eccDNA were extracted from genomic regions enriched in these circle-supporting reads. The aligned reads were visualized using IGV version 2.17.2.

Total RNAseq

Total RNA was extracted from frozen mouse tissue or frozen patient tissue using the RNeasy Micro kit (Quagen #74004). Total RNA samples were submitted to the Functional Genomic Center Zurich (FGCZ). RNAseq was performed using the Illumina Novaseq 6000 platform. Differential expression of genes and pathway analysis were performed by the FGCZ SUSHI platform. To perform GSEA analysis on the transcriptomic data, the datasets from WT, *Mcl1*$^{\Delta hep}$, and *Mcl1*$^{\Delta hep}$ *Sting1*$^{-/-}$ mice were imported to GSEA (4.3.3) Desktop Application and analyzed.

Graphical illustrations

Graphical illustrations shown in the main figures (Figure 2A, 3A, 4E and 4H left panel) were created with BioRender.com.

### Supplementary Tables

| **sample_id** | **chr_circle** | **start_circle** | **end_circle** | **chr_gene** | **start_gene** | **end_gene** | **gene_symbol** |
|---|---|---|---|---|---|---|---|
| WT1 | chr1 | 88139927 | 88198043 | chr1 | 88154708 | 88190011 | *Mroh2a* |
| WT1 | chr10 | 38696225 | 38697993 | chr10 | 38696537 | 38697775 | *Rfpl4b* |
| WT1 | chr17 | 13093453 | 13097236 | chr17 | 13094921 | 13096864 | *Mrgprh* |
| WT1 | chr17 | 13433078 | 13458279 | chr17 | 13440075 | 13446545 | *Smok2a* |
| WT1 | chr17 | 13433078 | 13458279 | chr17 | 13449144 | 13456071 | *Smok2b* |
| WT2 | chr1 | 88138955 | 88184803 | chr1 | 88139681 | 88146719 | *Ugt1a1* |
| WT2 | chr4 | 136332370 | 136356223 | chr4 | 136337748 | 136340537 | *Tex46* |
| WT2 | chr5 | 24639608 | 24655928 | chr5 | 24643438 | 24650285 | *Fastk* |
| WT2 | chr5 | 24639608 | 24655928 | chr5 | 24650456 | 24652852 | *Tmub1* |
| WT2 | chr5 | 25427846 | 25428588 | chr5 | 25427846 | 25428151 | *E130116L18Rik* |
| WT2 | chr8 | 19297542 | 19301227 | chr8 | 19297596 | 19300844 | *Defb5* |
| WT2 | chr8 | 115144223 | 115154275 | chr8 | 115144826 | 115152586 | *Clec3a* |
| WT2 | chr9 | 108736926 | 108752150 | chr9 | 108743687 | 108744631 | *Tmem89* |
| WT2 | chr17 | 13436747 | 13458908 | chr17 | 13440075 | 13446545 | *Smok2a* |
| WT2 | chr17 | 13436747 | 13458908 | chr17 | 13449144 | 13456071 | *Smok2b* |
| WT2 | chr19 | 5777890 | 5780417 | chr19 | 5778115 | 5779681 | *Fam89b* |
| WT2 | chr19 | 47139224 | 47155304 | chr19 | 47140048 | 47146203 | *Calhm3* |
| WT3 | chr1 | 88139117 | 88173315 | chr1 | 88139681 | 88146719 | *Ugt1a1* |
| WT3 | chr8 | 93624612 | 93631124 | chr8 | 93628035 | 93629039 | *Capns2* |
| WT3 | chr19 | 47137404 | 47150438 | chr19 | 47140048 | 47146203 | *Calhm3* |
| MCL1 | chr2 | 24859577 | 24866010 | chr2 | 24864129 | 24865110 | *Mrpl41* |
| MCL1 | chr2 | 27326957 | 27348407 | chr2 | 27332336 | 27334233 | *Brd3os* |
| MCL1 | chr4 | 132028593 | 132031030 | chr4 | 132029508 | 132030696 | *Rab42* |
| MCL1 | chr6 | 38085357 | 38093346 | chr6 | 38086190 | 38092744 | *Tmem213* |
| MCL1 | chr7 | 126969975 | 126975384 | chr7 | 126971709 | 126975222 | *Zfp747* |
| MCL1 | chr8 | 111782961 | 111785535 | chr8 | 111782971 | 111784296 | *Exosc6* |
| MCL1 | chr10 | 75605971 | 75609940 | chr10 | 75607064 | 75609248 | *Ddt* |
| MCL1 | chr13 | 23940875 | 23945433 | chr13 | 23940996 | 23941392 | *H4c2* |
| MCL1 | chr13 | 23940875 | 23945433 | chr13 | 23944675 | 23945213 | *H4c1* |
| MCL1 | chr17 | 27781671 | 27800171 | chr17 | 27782547 | 27784730 | *Smim29* |
| MCL1 | chrX | 94600343 | 94602643 | chrX | 94601710 | 94602409 | *Pfn5* |
| MCL2 | chr2 | 152403217 | 152412052 | chr2 | 152404897 | 152411458 | *Gm17416* |
| MCL2 | chr10 | 128042950 | 128048431 | chr10 | 128045777 | 128047676 | *Spryd4* |
| MCL2 | chr17 | 13442937 | 13458005 | chr17 | 13449144 | 13456071 | *Smok2b* |
| MCL2 | chr2 | 25388004 | 25392108 | chr2 | 25388663 | 25391731 | *C8g* |
| MCL2 | chr4 | 128998668 | 129007664 | chr4 | 128999341 | 129005419 | *Tmem54* |
| MCL2 | chr5 | 115435039 | 115441508 | chr5 | 115435169 | 115439058 | *Dynll1* |
| MCL2 | chr6 | 131662729 | 131666334 | chr6 | 131663495 | 131664458 | *Tas2r105* |
| MCL2 | chr7 | 126612862 | 126616695 | chr7 | 126614205 | 126616524 | *Pagr1a* |
| MCL2 | chr8 | 71915559 | 71919182 | chr8 | 71917603 | 71918391 | *Mrpl34* |
| MCL2 | chr9 | 108215480 | 108218880 | chr9 | 108216102 | 108217542 | *Gpx1* |
| MCL2 | chr10 | 82700868 | 82704799 | chr10 | 82702460 | 82703764 | *Eid3* |

| | | | | | | | |
|---|---|---|---|---|---|---|---|
| MCL2 | chr11 | 72101167 | 72112390 | chr11 | 72102550 | 72106418 | *Med31* |
| MCL2 | chr11 | 72101167 | 72112390 | chr11 | 72105964 | 72109270 | *4930563E22Rik* |
| MCL2 | chr11 | 120652816 | 120655283 | chr11 | 120653620 | 120655245 | *Hmga1b* |
| MCL3 | chr1 | 88138901 | 88184406 | chr1 | 88139681 | 88146719 | *Ugt1a1* |
| MCL3 | chr1 | 173100266 | 173106057 | chr1 | 173101949 | 173105344 | *Mptx2* |
| MCL3 | chr2 | 18000951 | 18002248 | chr2 | 18001228 | 18001769 | *H2al2a* |
| MCL3 | chr2 | 152577438 | 152580878 | chr2 | 152578171 | 152579330 | *Id1* |
| MCL3 | chr4 | 3870655 | 3874326 | chr4 | 3870657 | 3872105 | *Mos* |
| MCL3 | chr4 | 88439944 | 88445775 | chr4 | 88440262 | 88441011 | *Ifnb1* |
| MCL3 | chr4 | 118772679 | 118777852 | chr4 | 118773721 | 118774662 | *Or10ak8* |
| MCL3 | chr4 | 138694111 | 138700136 | chr4 | 138694423 | 138695676 | *Rnf186* |
| MCL3 | chr6 | 40470035 | 40475290 | chr6 | 40470463 | 40471475 | *Tas2r108* |
| MCL3 | chr6 | 96141391 | 96143923 | chr6 | 96141478 | 96143224 | *Nup50l* |
| MCL3 | chr6 | 132818659 | 132825642 | chr6 | 132824105 | 132825106 | *Tas2r123* |
| MCL3 | chr7 | 104382129 | 104386668 | chr7 | 104382592 | 104383557 | *Or52n1* |
| MCL3 | chr7 | 141710131 | 141711865 | chr7 | 141710191 | 141710850 | *Gm29735* |
| MCL3 | chr8 | 19243910 | 19247174 | chr8 | 19244373 | 19245228 | *Defb14* |
| MCL3 | chr11 | 99515502 | 99519621 | chr11 | 99518050 | 99519064 | *Krtap4-1* |
| MCL3 | chr13 | 110487329 | 110493929 | chr13 | 110489150 | 110493733 | *Gapt* |
| MCL3 | chr13 | 120776702 | 120780412 | chr13 | 120778414 | 120779714 | *Tcstv3* |
| MCL3 | chr15 | 98377329 | 98383171 | chr15 | 98378281 | 98380602 | *Lalba* |
| MCL3 | chr17 | 14168478 | 14172304 | chr17 | 14168635 | 14170940 | *Gm7168* |
| MCL3 | chr18 | 36856558 | 36859977 | chr18 | 36858120 | 36859851 | *Cd14* |
| MCL3 | chrX | 72271872 | 72274647 | chrX | 72271897 | 72274401 | *F8a* |

Table S1 – List of full genes present in eccDNA

| **Antibody** | **Company** | **Cat. #** | **Dilution** |
|---|---|---|---|
| Rabbit anti-MCL1 | abcam | ab32087 | 1:1000 |
| Mouse anti-β-Catenin | BD | 610153 | 1:400 |
| Mouse anti-γH2AX | BioLegend | 613410 | 1:100 |
| Rat anti-CldU | abcam | ab6326 | 1:400 |
| Mouse anti-IdU | BD | 347580 | 1:80 |
| Mouse anti-ssDNA | DSHB | 10805144 | 1:100 |
| Rabbit anti-Ki67 | abcam | ab16667 | 1:100 |
| Rabbit anti-Cl. Caspase 3 | Cell Signaling | 9664S | 1:100 |
| Rat anti-F4/80 | Biomedicals AG | T-2006 | 1:50 |
| Rabbit anti-γH2AX | Novus | NB100-384 | 1:200 |
| Rat anti-B220 | Pharmingen | 553084 | 1:4000 |
| Rat anti-Ly6G | Pharmingen | 551459 | 1:600 |
| Rabbit anti-CD3 | Thermo Fischer | MA1-90582 | 1:300 |
| Rabbit anti-cGAS | Cell Signaling | 31659S | 1:100 |
| Rabbit anti-STING | proteintech | 19851-I-AP | 1:1000 |

Table S2 - List of antibodies for IHC and IF

| **Antibody** | **Company** | **Cat. #** | **Dilution** |
|---|---|---|---|
| Rabbit anti- MCL1 | Rockland | 600-401-394 | 1:1000 |
| Rabbit anti-cGAS | Cell Signaling | 31659S | 1:1000 |
| Rabbit anti-STING | proteintech | 19851-I-AP | 1:1000 |
| Rabbit anti-GAPDH | Cell Signaling | 5174S | 1:1000 |
| Rabbit anti-p-p65 | Cell Signaling | 3033L | 1:1000 |
| Rabbit anti-p65 | Cell Signaling | 3034S | 1:1000 |
| Rabbit anti-CD45 | Cell Signaling | 72787S | 1:1000 |
| Rabbit anti-H3 | Cell Signaling | 4499S | 1:1000 |
| Rabbit anti-Albumin | Cell Signaling | 4929S | 1:1000 |
| Rabbit anti-p-IRF3 | Cell Signaling | 4947S | 1:1000 |

Table S3 - List of antibodies for Western blot

| **Antibody** | **Company** | **Cat. #** | **Dilution** |
|---|---|---|---|
| CD11b-FITC | BioLegend | 101205 | 1:100 |
| CD11c-V450 | BioLegend | 117329 | 1:100 |
| F4/80-PerCP | BioLegend | 123125 | 1:100 |

Table S4 - List of antibodies for flow cytometry

| Name | Sequence | Supplier |
|---|---|---|
| *Cgas*-F | CAGGCGTCTTCTTCAGTCATAC | Integrated DNA Technologies |
| *Cgas*-R | CTGGTCTACAGAGGGAGTTACA | Integrated DNA Technologies |
| *Sting1*-F | ACATTCCCTGCTCTCTGTTG | Integrated DNA Technologies |
| *Sting1*-R | TCCCTCTCGCCATCTACTT | Integrated DNA Technologies |

Table S5 – List of primers for qPCR

### References


1. Charni-Natan, M., Goldstein, I. Protocol for Primary Mouse Hepatocyte Isolation. *STAR Protoc*. **1**, 100086 (2020).
2. Toda, G., Yamauchi, T., Kadowaki, T., Ueki, K. Preparation and culture of bone marrow-derived macrophages from mice for functional analysis. *STAR Protoc*. **2**, 100246 (2021).
3. Toufektchan, E., Maciejowski, J. Purification of micronuclei from cultured cells by flow cytometry. *STAR Protoc*. **2**, 100378 (2021).
4. Wang, Y., Wang, M., Zhang, Y. Purification, full-length sequencing and genomic origin mapping of eccDNA. *Nat Protoc*. **18**, 683-99 (2023).
5. Henssen, A., MacArthur, I., Koche, R., H, D.-G. Purification and Sequencing of Large Circular DNA from Human Cells. *Protocol Exchange*. (2019).
